# A simple mechanism for epigenetic inheritance of silent chromatin

**DOI:** 10.1101/2023.07.18.549577

**Authors:** Andy H. Yuan, Danesh Moazed

## Abstract

Mechanisms enabling genetically identical cells to differentially regulate gene expression are complex and central to organismal development and evolution. While gene silencing pathways involving sequence-specific recruitment of histone-modifying enzymes are prevalent in nature, examples of sequence-independent heritable gene silencing are scarce. Studies of *Schizosaccharomyces pombe* indicate that sequence-independent propagation of heterochromatin can occur but requires numerous multisubunit protein complexes and their various activities. Such complexity has precluded a coherent articulation of the minimal requirements for heritable gene silencing by conventional approaches. Here, we take an unconventional approach to defining these requirements by engineering sequence-independent silent chromatin inheritance in *Saccharomyces cerevisiae*. The memory-conferring mechanism is remarkably simple and requires only two proteins, one that recognizes histone H3 methylation and deacetylates histone H4, and another that recognizes unmodified H4 and catalyzes H3 methylation. These bilingual “read-write” proteins form an interdependent positive feedback loop capable of transmitting sequence-independent silent information over multiple generations.

## Introduction

Heterochromatin preserves genome stability and downregulates gene expression. Histone H3 lysine 9 methylation (H3K9me) is a hallmark of heterochromatin in mammals, plants, and fungi including the fission yeast *Schizosaccharomyces pombe*. Heterochromatin-mediated gene silencing in *S. pombe* cells is heritable and couples RNA interference (RNAi) with the recognition and local propagation of H3K9me at centromeres and silent mating type cassettes^1–3^. Locus specificity is, in part, conferred by short interfering RNA (siRNA) sequence complementarity to nascent transcripts generated at target loci^3–5^. In addition to RNAi, *S. pombe* cells also employ DNA sequence-dependent mechanisms for gene silencing, whereby transcription factors bind specific DNA sequences and recruit heterochromatin-associated factors to chromatin via protein-protein interactions^6–8^.

The budding yeast *Saccharomyces cerevisiae*, which naturally lacks H3K9me and RNAi, assembles less complex heterochromatin at silent mating type loci via composite DNA sequences known as silencers^9^. Silencers feature binding sites for essential transcription factors, which recruit proteins of the silent information regulator (SIR) complex to regions flanking genes specifying mating type^10, 11^. The SIR complex is composed of Sir2, the founding member of the sirtuin-family of nicotinamide adenine dinucleotide (NAD^+^)-dependent deacetylases, Sir3, and Sir4. Sir2 deacetylates histone H4 lysine 16 (H4K16), and Sir3 recognizes unmodified H4K16- containing nucleosomes^12–14^. As Sir3 associates with Sir2 via Sir4^20^, the SIR complex propagates a hypoacetylated chromatin state in *cis* by physically spreading from silencers over mating type-specifying genes^15^.

Studies of heterochromatin in *S. pombe* cells revealed that silent chromatin states can propagate over multiple generations independently of underlying DNA sequence^16, 17^. Establishment and maintenance of these states require numerous factors, several enzymatic activities, and the absence of a crucial silencing antagonist^16–19^. In contrast, SIR complex-mediated gene silencing in *S. cerevisiae* cells requires the presence of silencer DNA at all times^20, 21^.

We reasoned that incorporating H3K9me into *S. cerevisiae* chromatin would enable us to test principles of heterochromatin-mediated gene silencing in a controllable environment ∼300 million years removed from the complexities of natural H3K9me-dependent processes. This bottom-up approach enabled us to engineer a reduced-complexity, heritable domain of silent chromatin, where silent information transfer over multiple generations occurs independently of DNA sequence and requires only two chimeric proteins.

## Results

### Introducing H3K9me to *S. cerevisiae* cells

We sought to deposit H3K9me at a specific locus in *S. cerevisiae* cells by adopting an experimental scheme that has been used to study inducible ectopic heterochromatin domains in *S. pombe* cells^16–18^. Specifically, we replaced the normally silent mating type locus *HMR* – including silencers *E* and *I* – with ten tandem tet operators (*tetO*_10X_) positioned upstream of an *ADE2* reporter gene in *ade2-1*Δ cells (Figure 1A). Cells in which *ADE2* is silent accumulate pigment and form red colonies on medium containing limiting adenine, whereas cells in which *ADE2* is active do not accumulate pigment and form white colonies. We then designed a chimeric protein consisting of the bacterial tetracycline repressor TetR fused to the catalytic SET (Suppressor of variegation, Enhancer of Zeste, Trithorax) domain of human Suppressor of Variegation 3-9 Homolog 2 (SUV39H2), an H3K9 methyltransferase (Figure 1A). We refer to this protein as TetR-SET, which contains a nuclear localization signal (NLS) and a V5 epitope tag between TetR and the SET domain (Figure S1A). Production of TetR-SET in reporter-containing cells resulted in local di- and tri-methylation of H3K9 at *tetO*_10X_ (Figures 1B and S1B). We took advantage of the reversible binding properties of TetR by treating cells with anhydrotetracycline (aTc) and measured the level of H3K9me2 under conditions where TetR- SET is no longer bound adjacent to *ADE2* (Figure 1A). Our results showed that the addition of aTc induced rapid TetR-SET untethering within 30 minutes (Figure S1C) and rendered H3K9me2 undetectable after six hours (Figure 1C).

**Figure 1.**
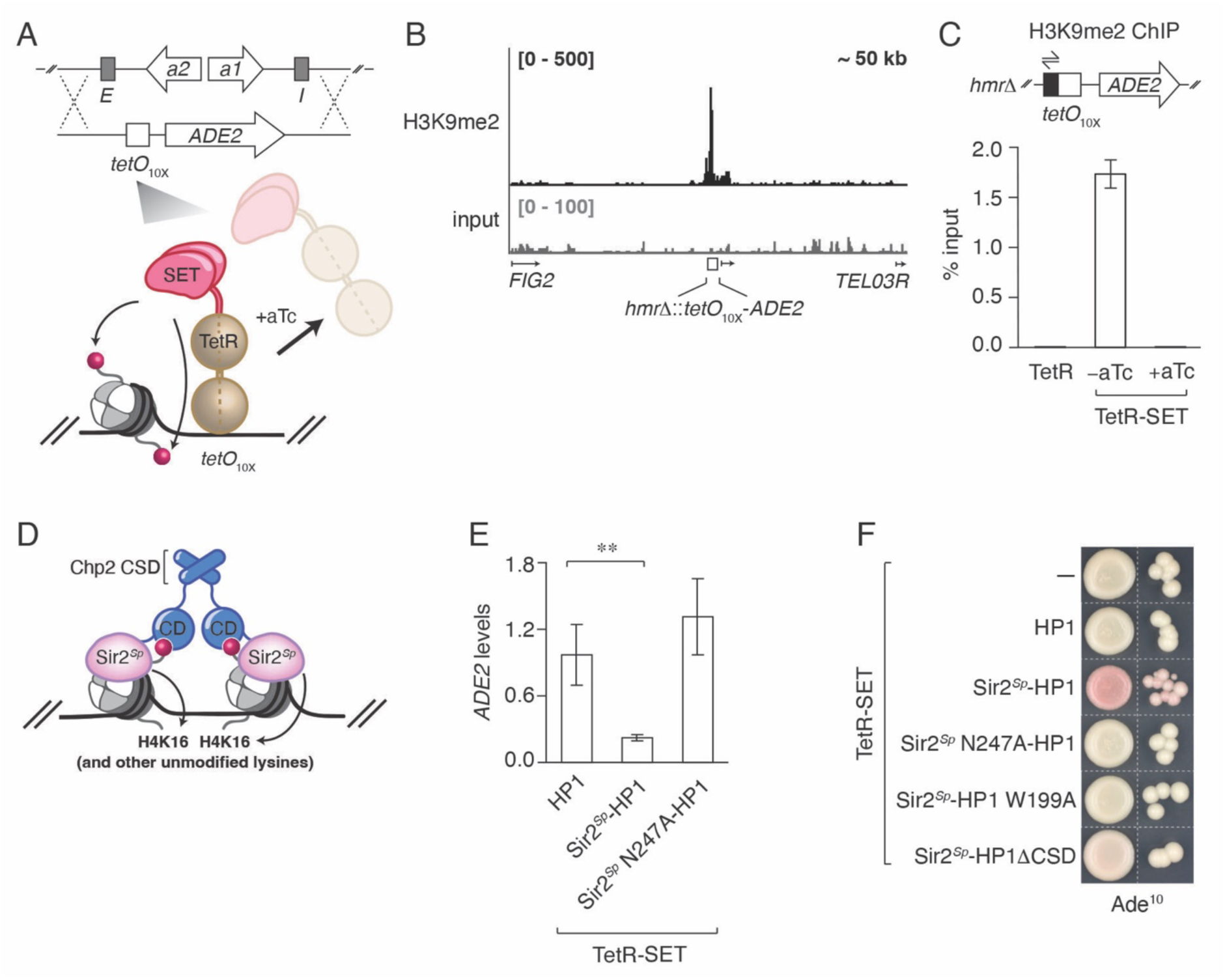
Establishment of H3K9me-dependent gene repression in *S. cerevisiae* cells. **(A)** Cartoon schematic of how *HMR* was replaced by a *tetO*_10X_-*ADE2* reporter via homologous recombination and how DNA sequence-specific recruitment of TetR-SET – consisting of bacterial TetR fused to the catalytic SET domain of human SUV39H2 – results in continuous local H3K9me (red spheres). TetR-SET is depicted as a dimer bound to a single *tet* operator. H3K9me ceases in the presence of anhydrotetracycline (aTc), which untethers TetR-SET from the *tet* operator array. **(B)** ChIP-seq profile of H3K9me2 at the *tetO*_10X_-*ADE2* reporter in cells producing TetR-SET. Data ranges (in counts per million) are indicated in brackets. The region visualized encompasses ∼50 kb of the right arm of chromosome III. The H3K9me2 profile is shown in red, and the input DNA profile is shown in gray. The position of *tetO*_10X_ is marked by a white square, and DNA features including select genes are indicated by arrows. **(C)** ChIP-qPCR analysis of H3K9me2 levels at the *tetO*_10X_-*ADE2* reporter in cells producing TetR or TetR-SET. Primers (half-headed arrows) bind to unique DNA sequences (black rectangle) adjacent to *tetO*_10X_ (white square). TetR-SET-producing cells were cultured in the presence or absence of aTc for six hours. Values represent the averages and standard deviations of three biological replicates. The level of H3K9me2 at the reporter locus in aTc-treated cells approached the limit of detection. **(D)** Cartoon depiction of Sir2*^Sp^*-HP1 - consisting of full-length *S. pombe* Sir2 fused to the chromodomain (CD), hinge region, and chromo shadow domain (CSD) of *S. pombe* Chp2 - bound as a dimer to a dinucleosome containing two methylated H3K9 residues (red spheres). Arrows are drawn to indicate Sir2*^Sp^*-mediated deacetylation of H4K16 and other lysines. **(E)** RT-qPCR analysis of *ADE2* mRNA levels in cells producing TetR-SET and either HP1, Sir2*^Sp^*-HP1, or a Sir2*^Sp^*-HP1 variant lacking histone deacetylase activity (Sir2*^Sp^* N247A-HP1). Values represent *ADE2* levels (normalized to *ACT1*) in the indicated cells relative to *ADE2* levels in reporter gene-containing cells producing TetR alone. Averages and standard deviations of three biological replicates are shown. Significance was assessed by a two-tailed Student’s t-test (*p* ≤ 0.01). **(F)** Phenotypes of *tetO*_10X_-*ADE2* reporter-containing cells producing TetR-SET and either HP1, Sir2*^Sp^*-HP1, Sir2*^Sp^* N247A-HP1, a Sir2*^Sp^*-HP1 variant containing a mutant chromodomain incapable of methyllysine recognition (Sir2*^Sp^*-HP1 W199A), or a monomeric Sir2*^Sp^*-HP1 variant lacking its chromo shadow domain (Sir2*^Sp^*-HP1ΔCSD). HP1, Sir2*^Sp^*-HP1, and Sir2*^Sp^*-HP1 variants were produced from pRS315, a low-copy plasmid encoding *LEU2*. Cells were spotted on leucine-lacking medium containing limiting adenine (10 μg/ml; Ade^10^). White dashed lines indicate sites of cropping between spots of cells grown and photographed on the same plate.

H3K9me recruits heterochromatin protein 1 (HP1)-family proteins to silent chromatin. HP1 proteins feature an H3K9me-recognizing chromodomain (CD) linked to a chromo shadow domain (CSD)^22^. The CSD mediates HP1 protein homodimerization as well as the direct or indirect recruitment of myriad effector proteins, including histone deacetylases (HDACs)^23–26^. As histone deacetylation is a conserved feature of heterochromatin-mediated gene silencing^12, 23, 24, 27^ and corepressor-mediated gene repression^28, 29^, we hypothesized that producing a single subunit HDAC fused to an HP1 protein would suffice to downregulate *ADE2* expression in TetR- SET-containing *S. cerevisiae* cells.

To test this hypothesis, we designed a chimeric protein consisting of *S. pombe* Sir2 (Sir2*^Sp^*) fused to an *S. pombe* HP1 protein (residues 170-380 of Chp2, encompassing the natural Chp2 chromodomain, hinge region, and chromo shadow domain) (Figure 1D). We refer to this protein as Sir2*^Sp^*-HP1, which contains an NLS and a Myc_3X_ epitope tag between Sir2*^Sp^* and HP1. Like its *S. cerevisiae* homolog Sir2*^Sc^*, Sir2*^Sp^* preferentially deacetylates H4K16^30^. As Sir3 recognizes unmodified H4K16-containing nucleosomes, all experiments described in this work were, unless otherwise noted, performed in *sir3*Δ cells to rule out contributions of the native *S. cerevisiae* SIR complex to reporter gene silencing.

Whereas production of HP1 in TetR-SET-containing cells had no effect on *ADE2* expression, production of Sir2*^Sp^*-HP1 resulted in decreased *ADE2* expression and increased colony pigmentation (Figures 1E and 1F). Downregulation of *ADE2* expression required the catalytic activity of Sir2*^Sp^*, a functional HP1 chromodomain, and HP1 dimerization (Figures 1E and 1F). A Sir2*^Sp^*-HP1 variant featuring the dimerization domain of bacteriophage λ CI in place of the Chp2 CSD behaved similarly to Sir2*^Sp^*-HP1 (Figure S1D). Reduced colony pigmentation was not attributable to differences in fusion protein levels (Figure S1E). We conclude that Sir2*^Sp^* HDAC activity is sufficient to downregulate *ADE2* expression, likely via the direct binding of Sir2*^Sp^*-HP1 to H3K9me-modified nucleosomes.

Three lines of evidence suggest that Sir2*^Sp^* acts independently of native *S. cerevisiae* silencing pathways: (i) the N-terminal domain of Sir2*^Sc^* – including its Sir4 interaction determinants – and the corresponding region of Sir2*^Sp^* lack sequence conservation and are structurally dissimilar (Figure S2A), (ii) unlike Sir2*^Sc^*-HP1, production of Sir2*^Sp^*-HP1 failed to rescue silencing in *sir2*Δ cells (Figure S2B), and (iii), unlike Sir2*^Sc^*, Sir2*^Sp^* had no discernable effect on silencing when fused to bacteriophage λ Cro and tethered to a TetR-independent reporter consisting of tandem *O_R_3* operators positioned upstream of the *ADE2* promoter (Figure S2C). We conclude – especially given that experiments were performed in *sir3*Δ cells – that Sir2*^Sp^*-HP1 functions independently of native *S. cerevisiae* silencing factors.

We asked if we could distinguish heterochromatin-mediated gene silencing, which is regional in nature, from gene repression, which can occur via the deacetylation of promoter-proximal nucleosomes^28, 29^. To do so, we modified our reporter by inserting another gene, *URA3* of *Kluyveromyces lactis* (*URA3^Kl^*), positioned upstream of *tetO*_10X_, with the *URA3^Kl^* promoter situated distal to the operator array (Figure 2A). As all cells carry a non-functional *ura3-1* allele, those in which *URA3^Kl^* is silent are resistant to 5-fluoroorotic acid (FOA) and those in which *URA3^Kl^* is active are not. If TetR-SET and Sir2*^Sp^*-HP1 repress *ADE2* expression primarily by modifying nucleosomes adjacent to *tetO*_10X_, then cells producing these two proteins alone are predicted to be sensitive to FOA, which was indeed the case (Figure 2A). We reasoned that *URA3^Kl^* silencing would require the propagation of H3K9me in *cis* over the *URA3^Kl^* gene body and promoter.

**Figure 2.**
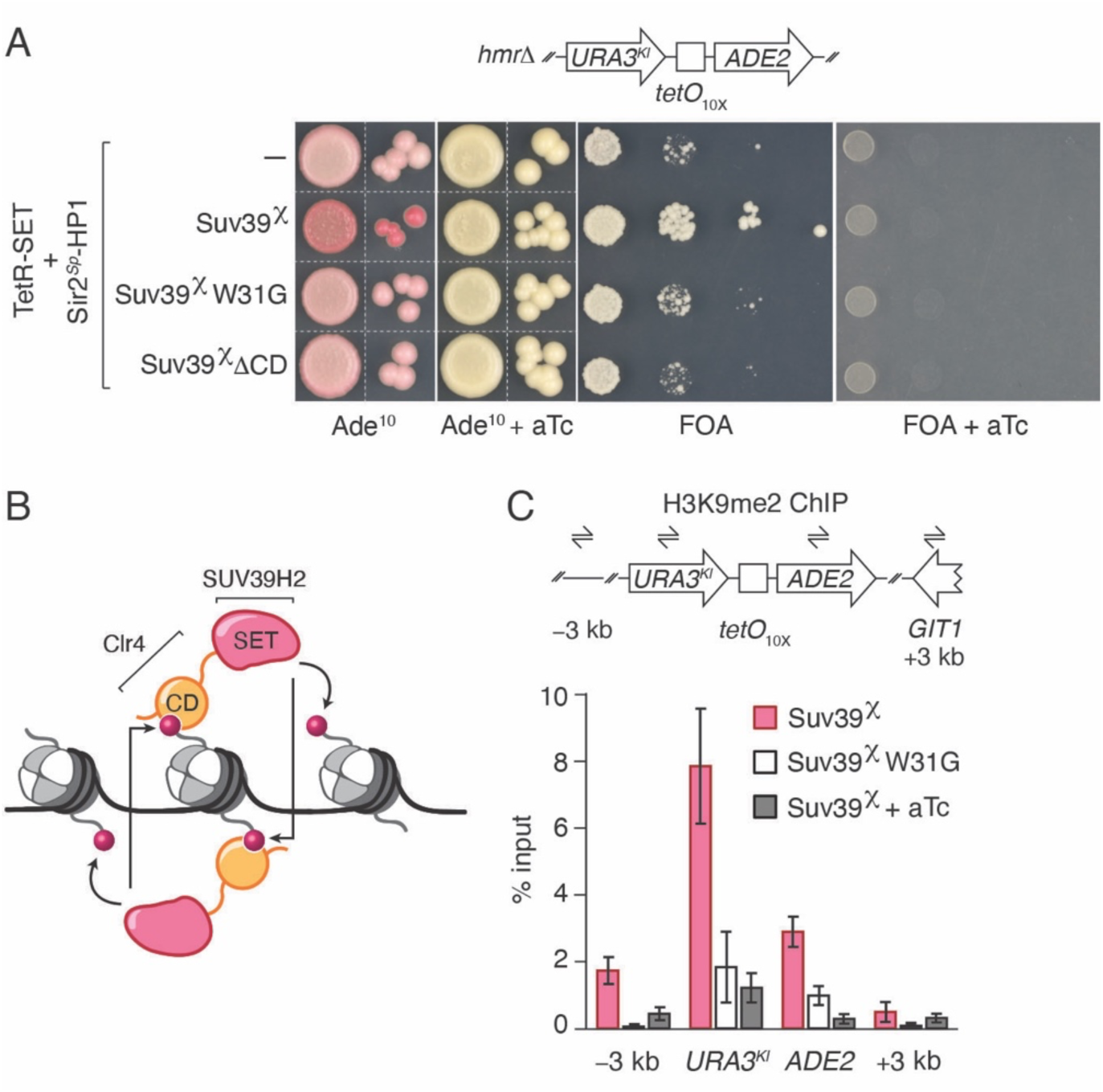
An H3K9me read-write protein allows for distinction between local gene repression and regional gene silencing in *S. cerevisiae* cells. **(A)** Schematic of the modified reporter and phenotypes of reporter-containing cells producing either TetR-SET and Sir2*^Sp^*-HP1 alone (row one), or TetR-SET and Sir2*^Sp^*-HP1 in combination with Suv39^χ^, a Suv39^χ^ variant containing a mutant chromodomain incapable of methyllysine recognition (Suv39^χ^ W31G), or a Suv39^χ^ variant lacking its chromodomain (Suv39^c^ΔCD) (rows two through four). Suv39^χ^ and Suv39^χ^ variants were encoded on pRS315. Cells were spotted on leucine-lacking medium containing the indicated compounds. White dashed lines indicate sites of cropping between spots of cells grown and photographed on the same plate. **(B)** Cartoon depiction of how Suv39^χ^ − consisting of the chromodomain (CD) and hinge region of *S. pombe* Clr4 fused to the catalytic SET domain of human SUV39H2 – enables H3K9me spreading. Two Suv39^χ^ monomers that “read” a nucleosome containing two methylated H3K9 residues (red spheres) and “write” H3K9me on adjacent nucleosomes are illustrated. **(C)** ChIP-qPCR analysis of H3K9me2 levels at the *URA^Kl^*-*tetO*_10X_-*ADE2* reporter and sites located ∼3 kb away from *tetO*_10X_. Primer binding sites are indicated by half-headed arrows. Cells produce TetR-SET, Sir2*^Sp^*-HP1, and either Suv39^χ^ (pink) or Suv39^χ^ W31G (white). Suv39^χ^-producing cells were cultured in the presence (gray) or absence (pink) of aTc for six hours. Values represent the averages and standard deviations of three biological replicates.

SUV39H-family proteins are uniquely poised to support H3K9me propagation due to their ability to recognize (or “read”) and catalyze (or “write”) the same histone modification^31^. This read-write capability is the basis for a positive feedback loop proposed to enable spreading of H3K9me at target loci while ensuring the faithful transfer of silent information from parental to newly synthesized histones following DNA replication^16, 17, 32^. We therefore sought to test whether an enzyme capable of reading and writing H3K9me would permit H3K9me spreading and maintain reporter gene silencing without continuous DNA sequence-dependent recruitment of H3K9 methyltransferase activity.

To do so, we designed a chimeric H3K9me read-write protein consisting of the chromodomain and hinge region of the *S. pombe* H3K9 methyltransferase Clr4 (residues 1-191 of Clr4) fused to the SET domain of human SUV39H2 (Figure 2B) (the natural SET domain of Clr4 exhibited reduced function relative to that of SUV39H2 in *S. cerevisiae* cells). We refer to this protein as Suv39 chi(mera) (Suv39^χ^), which contains an NLS and a FLAG_3X_ epitope tag at its N-terminus. Cells producing TetR-SET, Sir2*^Sp^*-HP1, and Suv39^χ^ containing a functional chromodomain formed colonies exhibiting increased pigmentation and were resistant to FOA, suggesting that Suv39^χ^ had the capacity to read and write H3K9me (Figure 2A). In contrast, cells producing Suv39^χ^ lacking its chromodomain or a Suv39^χ^ variant harboring a missense substitution abolishing methyllysine recognition (residue W31G of Clr4) exhibited no change in colony pigmentation and were sensitive to FOA (Figure 2A). Reduced colony pigmentation and FOA-sensitivity were not attributable to differences in fusion protein levels (Figure S3A), and all requirements for Sir2*^Sp^*-HP1 function established previously for cells lacking Suv39^χ^ also applied to cells containing Suv39^χ^ (Figure S3B). However, cells grown in the presence of aTc formed unpigmented colonies and were sensitive to FOA (Figure 2A). Analysis of chromatin immunoprecipitates showed that H3K9me2 was present at *URA3^Kl^* and *ADE2* promoters and gene bodies (Figure S3C). Furthermore, the extent and local density of H3K9me2 was dependent on the read function of Suv39^χ^ (Figure 2C). Yet Suv39^χ^ could not maintain local H3K9me2 levels in the presence of aTc (Figure 2C), a result that we found puzzling.

### Sequence-independent heritable gene silencing in *S. cerevisiae* cells

We reasoned that the inability of Suv39^χ^ to maintain reporter gene silencing could be explained by the following logic: one chromodomain of Suv39^χ^ necessarily competes with two chromodomains of the Sir2*^Sp^*-HP1 dimer for association with H3K9me-modified nucleosomes. FOA-resistant cells must therefore produce Suv39^χ^ at sufficiently high levels to support local spreading. High levels of Suv39^χ^, however, can result in off-target H3K9me at sites other than the reporter locus (Figure S3D). Off-target H3K9me, in turn, redistributes Suv39^χ^ and Sir2*^Sp^*- HP1 genome-wide, effectively diluting both proteins and thereby rendering locus-specific reporter gene silencing unsustainable without continuous sequence-specific recruitment of TetR- SET.

Based on this logic, we hypothesized that sustainable locus-specific silencing might be achieved by transplanting the H3K9 methyltransferase activity of Suv39^χ^ to a naturally dimer-forming protein capable of recognizing a histone modification other than H3K9me. To test this hypothesis, we designed a chimeric protein composed of full-length Sir3 – featuring an unmodified H4K16-recognizing bromo-adjacent homology (BAH) domain and a winged helix (wH) homodimerization domain^13, 33^ – fused to the SET domain of human SUV39H2. We refer to this protein as Sir3-SET, which contains an NLS and FLAG_3X_ epitope tag between Sir3 and the SET domain. In the absence of aTc, cells producing Sir3-SET and Sir2*^Sp^*-HP1 accumulated more pigment and were more resistant to FOA than Suv39^χ^-containing cells (Figure 3A). These phenotypes were attributable to decreased reporter gene expression and reduced RNA polymerase II (Pol II) occupancy at the singular *ADE2* reporter gene promoter (Figures S4A and S4B). In the presence of aTc, however, only cells producing Sir3-SET accumulated pigment and remained resistant to FOA (Figure 3A). As Sir3 naturally interacts with Sir2-bound Sir4^34^, we deleted *SIR2* and *SIR4*, thereby rendering cells *sir2*Δ *sir3*Δ *sir4*Δ. We refer to cells lacking all three components of the SIR complex as *sir*^−^. Critically, production of Sir3-SET and Sir2*^Sp^*-HP1 in *sir*^−^ cells was sufficient to establish and maintain reporter gene silencing (Figure 3A). Silencing required a functional Sir3 BAH domain and Sir3 dimerization (Figure 3B). Reduced colony pigmentation and FOA-sensitivity were not attributable to loss of fusion protein production (Figure S4C). We conclude that Sir3-SET and Sir2*^Sp^*-HP1 together form an interdependent positive feedback loop capable of supporting DNA sequence-independent heritable gene silencing (Figure 3C).

**Figure 3.**
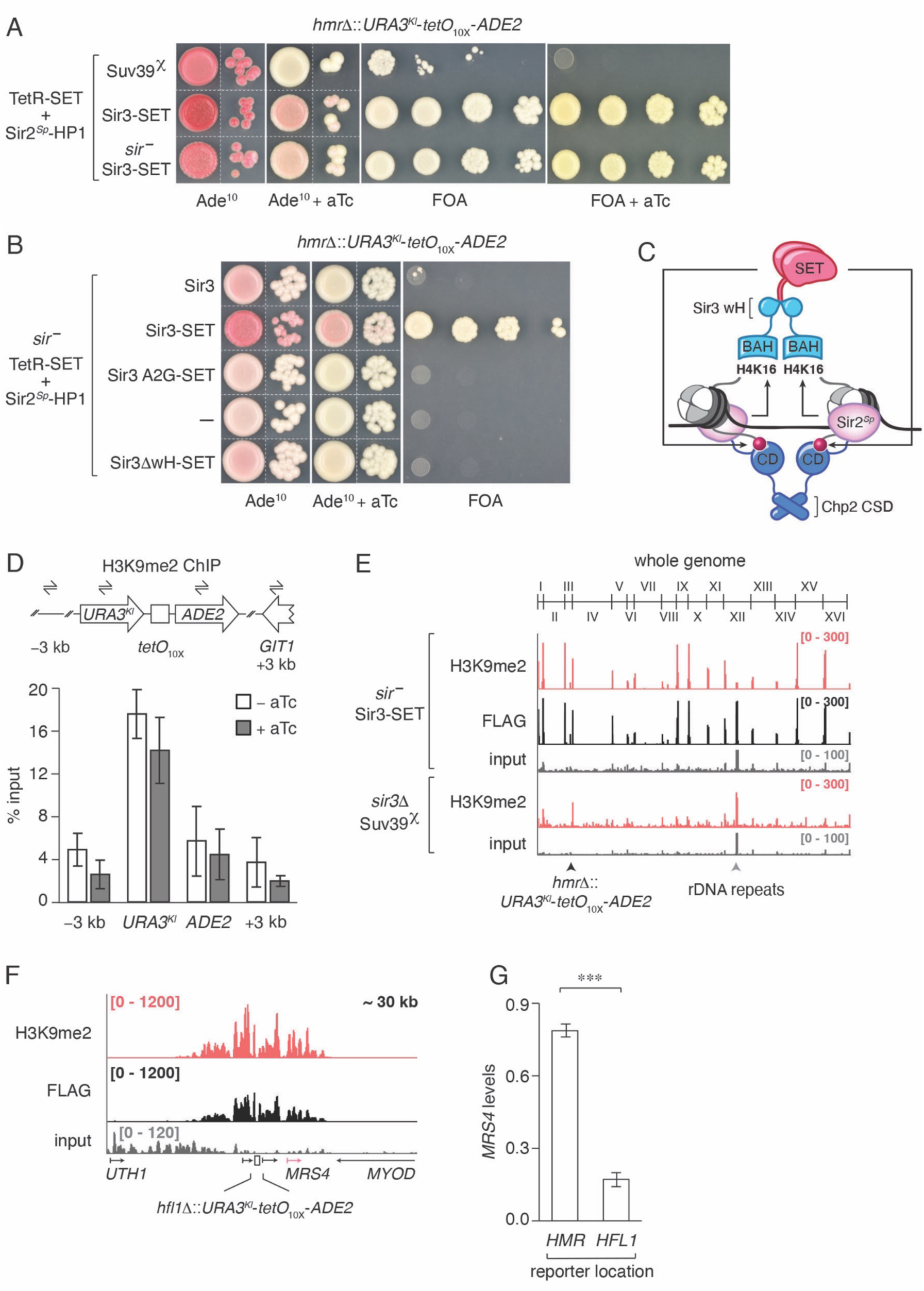
An interdependent positive feedback loop maintains DNA sequence-independent gene silencing in *S. cerevisiae* cells. **(A)** Phenotypes of *URA3^Kl^*-*tetO*_10X_-*ADE2* reporter-containing cells producing TetR-SET, Sir2*^Sp^*- HP1, and either Suv39^χ^ (row one) or Sir3-SET. Sir3-SET was produced in *sir3*Δ cells (row two) or *sir2*Δ *sir3*Δ *sir4*Δ (*sir*^−^) cells (row three). Suv39^χ^ and Sir3-SET were encoded on pRS315. Cells were spotted on leucine-lacking medium containing the indicated compounds. White dashed lines indicate sites of cropping between spots of cells grown and photographed on the same plate. **(B)** Phenotypes of *URA3^Kl^*-*tetO*_10X_-*ADE2* reporter-containing *sir*^−^ cells producing TetR-SET and Sir2*^Sp^*-HP1 alone, or TetR-SET and Sir2*^Sp^*-HP1 in combination with either Sir3, Sir3-SET, a Sir3- SET variant harboring a mutation that prevents Sir3 acetylation and disrupts bromo adjacent homology domain structure (Sir3 A2G-SET), or a monomeric Sir3-SET variant lacking its winged helix (wH) domain (Sir3ΔwH-SET). Sir3, Sir3-SET, and Sir3-SET variants were encoded on pRS315. Cells were spotted on leucine-lacking medium containing the indicated compounds. White dashed lines indicate sites of cropping between spots of cells grown and photographed on the same plate. **(C)** Cartoon depiction of the interdependent positive feedback loop formed by a pair of bilingual “read-write” proteins: Sir2*^Sp^*-HP1 and Sir3-SET. Sir3-SET is shown bound as a dimer to a dinucleosome containing two unmodified H4K16 residues. The Sir3 winged helix (wH) domain mediates Sir3-SET dimerization, and the Sir3 bromo adjacent homology (BAH) domain mediates the recognition of unmodified H4K16 by Sir3-SET. Sir3-SET methylates nearby H3K9 residues of the dinucleosome to which it is bound (as illustrated) as well as H3K9 residues of flanking nucleosomes (not illustrated). Two newly methylated H3K9 residues (red spheres), in turn, recruit the Sir2*^Sp^*-HP1 dimer (see Figure 1D). Sir2*^Sp^*-HP1 deacetylates nearby H4K16 residues (and other lysines) of the dinucleosome to which it is bound (as illustrated) in addition to H4K16 (and other lysines) of flanking nucleosomes (not illustrated). Two newly deacetylated H4K16 residues, in turn, recruit the Sir3-SET dimer. Arrows are drawn to indicate either Sir3- SET-mediated methylation of H3K9 or Sir2*^Sp^*-HP1-mediated deacetylation of H4K16 and other lysines. **(D)** ChIP-qPCR analysis of H3K9me2 levels at the *URA^Kl^*-*tetO*_10X_-*ADE2* reporter and sites located ∼3 kb away from *tetO*_10X_. Primer binding sites are indicated by half-headed arrows. Cells are *sir*^−^ and produce TetR-SET, Sir2*^Sp^*-HP1, and Sir3-SET. Cells were cultured for six hours in the presence (gray) or absence (white) of aTc. Values represent the averages and standard deviations of three biological replicates. **(E)** ChIP-seq profiles of H3K9me2 and Sir3-SET in *URA3^Kl^*-*tetO*_10X_-*ADE2* reporter-containing *sir*^−^ cells producing TetR-SET and Sir2*^Sp^*-HP1. Sir3-SET contains a FLAG_3X_ epitope tag. Data ranges (in counts per million) are indicated in brackets. The whole genome is visualized, with telomere junctions for all 16 chromosomes indicated on the hash-marked line. H3K9me2 profiles are shown in red, the Sir3-SET profile is shown in black, and input DNA profiles are shown in gray. The H3K9me2 profile of reporter-containing *sir3*Δ cells producing TetR-SET, Sir2*^Sp^*-HP1, and Suv39^χ^ is shown for comparison. Black and gray arrowheads point to the reporter locus and rDNA repeats, respectively. **(F)** ChIP-seq profiles of H3K9me2 and Sir3-SET in *sir*^−^ cells containing the *URA3^Kl^*-*tetO*_10X_- *ADE2* reporter in place of *HFL1*. In addition to Sir3-SET, which contains a FLAG_3X_ epitope tag, cells also produce TetR-SET and Sir2*^Sp^*-HP1. Data ranges (in counts per million) are indicated in brackets. The region visualized encompasses ∼30 kb of chromosome XI centered about *tetO*_10X_. The position of *tetO*_10X_ is indicated by a white rectangle, and select genes are indicated by arrows. *MRS4*, which is naturally positioned adjacent to *HFL1*, is highlighted in pink. **(G)** RT-qPCR analysis of *MRS4* mRNA levels in *sir*^−^ cells containing the *URA3^Kl^*-*tetO*_10X_-*ADE2* reporter in place of either *HMR* or *HFL1*. Cells produce TetR-SET, Sir2*^Sp^*-HP1, and Sir3-SET. Values represent *MRS4* levels (normalized to *ACT1*) in cells carrying the reporter in place of the indicated locus relative to *MRS4* levels in reporter-containing cells producing TetR alone. Averages and standard deviations of three biological replicates are shown. Significance was assessed by a two-tailed Student’s t-test (*p* ≤ 0.001).

Analysis of chromatin immunoprecipitates confirmed that H3K9me2 levels persist at reporter genes in interdependent positive feedback loop-containing *sir*^−^ cells after six hours of growth in the presence of aTc (Figure 3D). However, in addition to reporter genes, Sir3-SET and H3K9me2 also localized to telomeres (Figure 3E). *S. cerevisiae* telomeres consist of repetitive DNA sequence featuring binding sites for the essential transcription factor Rap1. Rap1 recruits the SIR complex to telomeres via protein-protein interactions with Sir4 and Sir3^10, 35^. As *sir*^−^ cells lack Sir4, we suspected that telomere localization resulted from Sir3-SET interacting with telomere-bound Rap1. To assess the potential contribution of DNA sequence – particularly that of the *HMR*-proximal telomere – to reporter gene silencing, we performed three tests.

First, to rule out contribution of DNA-bound Rap1 to reporter gene silencing, we deleted Rap1-interaction determinants of Sir3 (residues 456-479) from Sir3-SET^10^. *sir*^−^ cells producing Sir3Δ456-479-SET behaved similarly to those producing Sir3-SET (Figure S4D), indicating that neither establishment nor maintenance of reporter gene silencing requires an interaction between Sir3-SET and DNA-bound Rap1.

Second, to rule out contribution of other DNA sequences in the vicinity of *HMR* to reporter gene silencing, we relocated *tetO*_10X_ – along with flanking *URA3^Kl^*and *ADE2* reporter genes – from chromosome III to the non-essential, centromere-proximal, and normally euchromatic *HFL1* locus of chromosome XI. Analyses of colony color phenotypes and chromatin immunoprecipitates showed that cells producing Sir3-SET and Sir2*^Sp^*-HP1 silenced reporter genes – as well as a gene naturally flanking *HFL1* – and maintained silencing in the presence of aTc. These results indicate that reporter gene silencing does not require a specific chromatin context (Figures S4E, 3F, and 3G).

Third, to rule out any residual contribution of *tetO*_10X_ to reporter gene silencing in aTc-treated cells, we deleted the gene encoding TetR-SET (Figure S5). We performed this deletion in cells cultured in the presence of FOA. *URA^Kl^* silencing can therefore be maintained under selection in cells devoid of TetR-SET (Figure 4A). FOA-resistant cells lacking TetR-SET accumulated an intermediate level of pigment on medium containing FOA and limiting adenine (Figure 4A). *URA3^Kl^* and *ADE2* were therefore co-silenced under selection, consistent with the regional nature of heterochromatin. However, unlike cells containing TetR-SET, cells lacking TetR-SET formed colonies exhibiting little to no pigmentation on non-selective medium (Figure 4A). Based on these results, we conclude that interdependent positive feedback loop-mediated reporter gene silencing does not require specific DNA sequence and is intrinsically unstable without selection^17, 36^.

**Figure 4.**
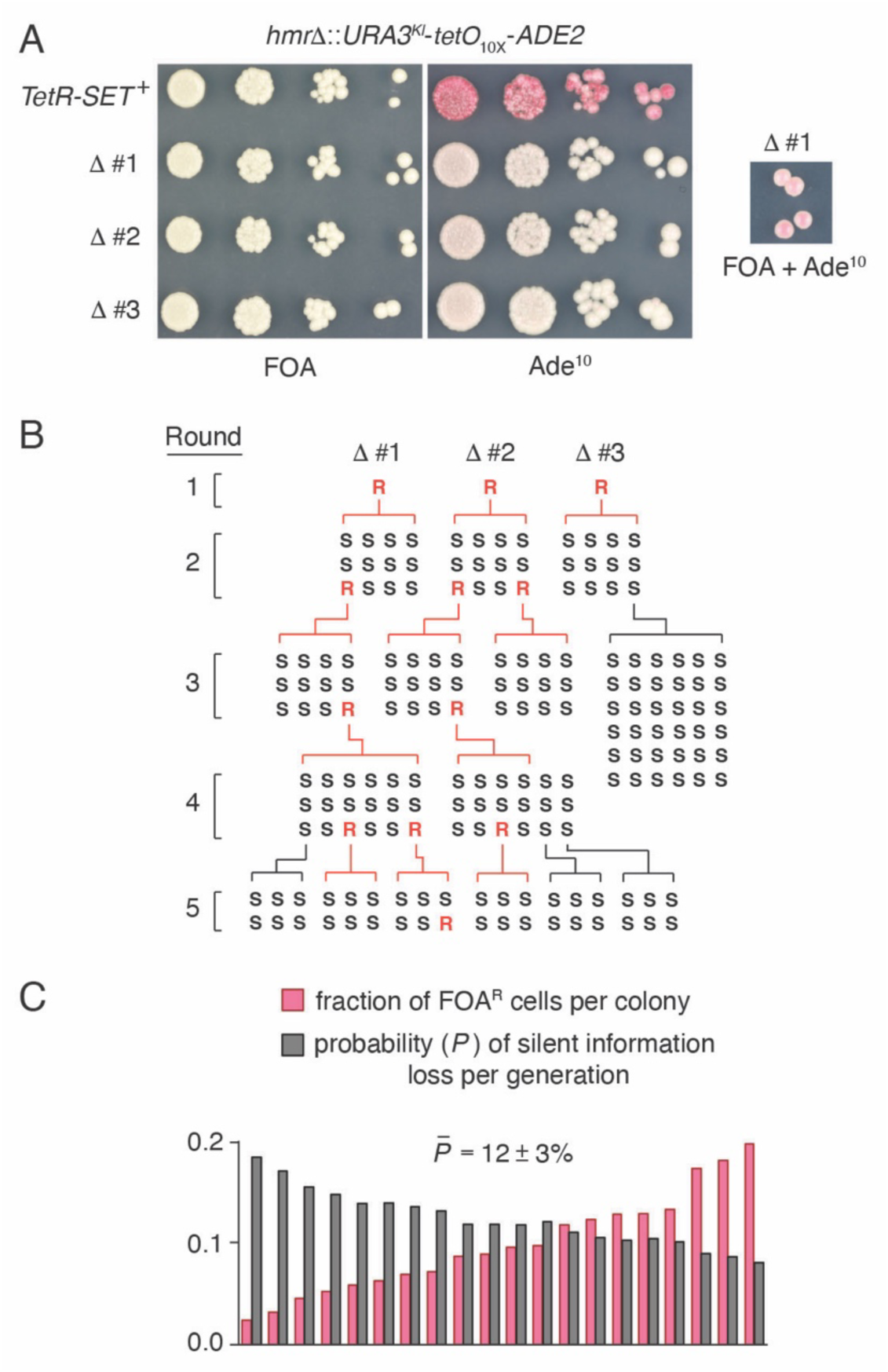
Reporter gene silencing is heritable but intrinsically unstable. **(A)** Phenotypes of *sir*^−^ cells producing Sir2*^Sp^*-HP1 and Sir3-SET in the presence or absence of TetR-SET-encoding DNA. Three strains lacking TetR-SET (Δ #1-3) isolated on and maintained in FOA-containing medium are shown. Cells were grown in the presence of FOA and spotted on medium containing FOA or Ade^10^. The phenotype of Δ #1 cells grown on medium containing FOA and Ade^10^ is indicative of *URA3^Kl^*and *ADE2* reporter gene co-silencing. **(B)** Silent information can propagate over multiple generations. Colonies – established by a Δ #1, Δ #2, or Δ #3 FOA-resistant founder cell – grown without selection either contain some fraction of FOA-resistant cells (red letter R) or contain FOA-sensitive cells only (black letter S). FOA-resistant daughter cells can be traced from round to round and emerge only from FOA- resistant mother cells, as evidenced by the Δ #1 lineage. Cells are unlikely to acquire resistance to FOA spontaneously, and once resistance to FOA is lost, it becomes irretrievable, as exemplified by the Δ #3 lineage. **(C)** For 20 Round 1 colonies grown on adenine-replete medium lacking FOA, the fraction of FOA-resistant (FOA^R^), pigment-accumulating cells per colony was determined (pink), and the probability (*P*) of silent information loss per generation was calculated (gray) (see Materials and Methods). The final calculated probability of silent information loss per generation represents the average and standard deviation of *P* for all 20 colonies.

Two possible scenarios can account for the apparent lack of pigment accumulation in cells devoid of TetR-SET: either (i) colonies contain homogeneous populations of cells that all gradually lose silent information during colony formation, or (ii) colonies contain heterogeneous populations of cells, a minority of which retain silent information, and a majority of which lose silent information^37^. To distinguish between these possibilities, we asked whether colonies – each established by an FOA-resistant founder cell cultured under selection – still contain FOA-resistant cells after growth on medium lacking FOA, and if so, whether FOA resistance was heritable.

To do so, we plated TetR-SET-lacking cells cultured in the presence of FOA on adenine- replete medium lacking FOA. Cells grown under adenine-replete conditions do not accumulate pigment and therefore form white colonies regardless of *ADE2* levels. We randomly picked colonies and patched them onto medium containing limiting adenine and FOA. *URA3^Kl^*and *ADE2* co-silencing allowed for the distinction between reporter gene silencing and genetic mutation based on whether a patch was pigmented or white. All of these colonies, which we refer to as Round 1 colonies, formed pigmented patches on FOA-containing medium (Figure 4B). Round 1 colonies patched onto selective medium were also restreaked on adenine-replete medium lacking FOA for a second round of colony formation without selection. Unlike Round 1 colonies, only 3 out of 36 Round 2 colonies – each established by a founder cell, the phenotype of which was initially unknown – formed pigmented patches on FOA-containing medium (Figure 4B). We patched and restreaked cells for three additional rounds and found that reporter gene silencing was detectable without selection through Round 5 (encompassing ∼15 days and ∼100 generations) (Figure 4B). We conclude that DNA sequence-independent silent information propagates in a minority of cells belonging to a heterogeneous population, and that cells retaining silent information undergo rapid extinction.

To quantify the (in)stability of reporter gene silencing, we calculated the probability that silent information-containing mother cells give rise to silent information-lacking daughter cells in a population undergoing exponential growth. To do so, we estimated the number of pigment- accumulating, FOA-resistant cells in each of 20 Round 1 colonies by resuspending and replating ∼250 cells onto selective and non-selective medium (Figure 4C). We used the fraction of FOA-resistant cells per colony to calculate the probability (*P*) of silent information loss per generation (see Materials and Methods). Assuming that differences in the growth rate and replicative lifespan of cells carrying either silent or active reporter genes are negligible during colony formation on adenine-replete medium lacking FOA, we conclude that the probability of silent information loss per generation is ∼12% (Figure 4C). The result of our calculation is surprisingly similar to measurements of ectopic heterochromatin stability in *S. pombe* cells that were based on colony sectoring phenotypes^17^. It is unlikely that our engineered *S. cerevisiae* cells transmit silent information as efficiently as natural *S. pombe* cells do. We therefore propose that previous estimates of stability were conservative and that protein factors and nucleic acid sequence elements – either characterized or awaiting discovery – likely reinforce silent information transfer in *S. pombe* cells.

### A requirement for native *S. cerevisiae* replisome-associated factors

To ask whether our system is regulated by any native *S. cerevisiae* processes, we performed a transposon mutagenesis screen in *sir*^−^ cells containing a *tetO*_10X_-*ADE2* reporter along with TetR-SET, Sir2*^Sp^*-HP1, and Sir3-SET. In total, we screened ∼25,000 transposon-containing mutants for altered colony pigmentation on medium lacking aTc and ∼12,500 mutants on medium containing aTc. We failed to isolate a single mutant exhibiting increased colony pigmentation in the presence of aTc – suggesting that *S. cerevisiae* cells naturally lack non-essential, H3K9me-specific silencing antagonists – but isolated 78 mutants exhibiting decreased colony pigmentation in the absence of aTc. We mapped transposon insertion sites for all 78 mutants, many of which were located at the promoter region of the gene encoding Sir3-SET or the reporter locus itself. Strikingly, however, the majority of remaining transposon insertions mapped to genes – most of which were identified at least twice and up to five times – involved in one of two processes: (i) NAD^+^ precursor metabolism, or (ii) DNA replication (Figure S6A). While we focused on the latter, we note that the availability of histone modifying enzyme cofactors – in this case, NAD^+^-dependent Sir2*^Sp^*-HP1 – can limit heterochromatin-mediated gene silencing^38^.

We deleted the coding sequence of five genes that were identified in our screen and encode replisome-associated proteins in either *sir3*Δ cells containing a *URA3^Kl^*-*tetO*_10X_-*ADE2* reporter, TetR-SET, Sir2*^Sp^*-HP1, and Sir3-SET, or *SIR^+^* cells containing an *ADE2* reporter gene situated between the natural *E* and *I* silencers of *HMR*. Deletion of four genes (*DPB4*, *MRC1*, *TOF1*, and *CTF4*) in interdependent positive feedback loop-containing cells resulted in reduced colony pigmentation and rendered cells sensitive to FOA, even in the absence of aTc (Figure S6B). In contrast, deletion of *ELG1*, the product of which is thought to play a pivotal role in natural *S. cerevisiae* gene silencing^39^, had more subtle effects. The same gene deletions had modest to no effects on colony pigmentation in *SIR^+^* cells (Figure S6C), suggesting that silencing defects were unlikely to be the result of genome-wide nucleosome depletion. Further characterization of cells lacking either *DPB4*, encoding a subunit of DNA polymerase ε (Pol ε) required for symmetric parental histone transfer to nascent DNA^40^, or *MRC1*, encoding an S phase checkpoint protein bridging Pol ε and the replisome helicase^41^, showed that mutant phenotypes were indicative of increased reporter gene expression (Figure S6D) and were not the result of reduced fusion protein levels (Figure S6E). These results suggest that interdependent positive feedback loop-containing cells are sensitive to asymmetric parental histone transfer and require *MRC1* to establish reporter gene silencing.

At first glance, it is surprising that the deletion of genes encoding replisome-associated factors impaired reporter gene silencing in the absence of aTc, a condition under which DNA sequence-dependent recruitment of H3K9 methyltransferase activity occurs continuously. In fact, replisome-associated mutations isolated in *S. pombe* cells have been shown to specifically affect the maintenance of heterochromatin, not its establishment^19^. We reason, however, that if mechanisms underlying heterochromatin-mediated gene silencing require time to reach and maintain critical densities of histone modifications at target loci, then idiosyncrasies of *S. pombe* and *S. cerevisiae* cell cycles may account for the sensitivity of interdependent positive feedback loop-containing cells to defects in chromatin replication^42–44^. Whereas *S. pombe* cells spend roughly 70% of each cell cycle in G2 phase, *S. cerevisiae* cells commit to mitosis shortly after completing faithful DNA synthesis. Histone H3 serine 10 (H3S10) phosphorylation evicts HP1 from chromatin during mitosis^45^, posing yet another obstacle to H3K9me-dependent silent information transfer. We propose that cells rely on two post-replicative windows of opportunity – once in G2 and again in G1 following H3S10 dephosphorylation – to both establish and maintain H3K9me-dependent heterochromatin. One of these windows is shut to *S. cerevisiae* cells, which may therefore lack the time required to establish reporter gene silencing in the absence of a fully functional replisome.

## Discussion

The prime objective of many branches of molecular biology is to reduce the complexity of otherwise staggeringly complex natural phenomena, a process traditionally falling under the jurisdiction of biochemistry and *in vitro* reconstitution. Our work demonstrates how reductionism can be applied *in vivo* to reveal the core requirements for histone modification-based memory of silent information. Specifically, we distilled the natural complexities of locus-specific, heritable gene silencing to their essential components: a pair of bilingual read-write proteins, each of which (i) binds cooperatively to nucleosomes, (ii) reads one of two histone modifications, and (iii) writes the modification distinct from what it reads, with one of the modifications being a specificity-determinant as well as a contributor to chromatin silencing. Interdependent positive feedback-mediated gene silencing (requiring only two chimeric proteins and two nucleosomal specificity-determinants - H3K9me and unmodified H4K16, in the case of *S. cerevisiae* cells described here), as opposed to natural *S. pombe* heterochromatin-mediated gene silencing (requiring numerous factors in addition to a “monolingual”, H3K9me-specific read-write protein), provides a simple mechanism by which cells can silence specific chromosomal loci over multiple generations independently of underlying DNA sequence.

Coupled feedback loops feature prominently in heritable gene silencing systems throughout Eukarya. For example, H3K9me and RNAi are coupled in fungi and plants^46^, histone H3 lysine 27 methylation and histone H2A lysine 119 ubiquitination are coupled in mammals^47^, and DNA methylation is coupled to H3K9me in fungi, plants, and mammals^48–50^. Operating principles of the sequence-independent heritable gene silencing system described here promise to inform studies of these and other potentially memory-conferring cellular processes.

## Acknowledgments

We thank N. Maier for advice on ChIP-seq data processing, T. Bartlett for advice on calculating the probability of silent information loss, and N. Craig for the gift of pSG36. We are indebted to A. Hochschild and F. Winston for invaluable discussions and comments on the manuscript. This work was supported by an NIH K99 Pathway to Independence Award (K99 GM137045-01 to A.H.Y.), an NIH postdoctoral fellowship (F32 GM131438-01 to A.H.Y.), and an NIH grant (R01GM072805 to D.M.). D.M. is an investigator of the Howard Hughes Medical Institute.

## Author contributions

D.M. and A.H.Y conceived of the experimental approach. A.H.Y. designed and performed experiments. A.H.Y. analyzed data. A.H.Y. and D.M. wrote the paper.

## Materials and Methods

### *Saccharomyces cerevisiae* cell growth, genetic manipulation, and plate-based phenotypic analysis

*S. cerevisiae* cells derived from strain W303-1a were grown in YPAD (1% yeast extract, 2% peptone, and 2% glucose supplemented with 50 μg/ml tryptophan and 50 μg/ml adenine) or synthetic medium in the presence or absence of 10 μM anhydrotetracycline (aTc) at 30°C.

Genes were integrated onto chromosomes in two steps. First, a target locus (either *ade2- 1*, *LYS2*, or *CAN1*) was replaced with a *URA3^Kl^*- and *KanMX*-encoding cassette conferring sensitivity to 1 mg/ml 5-fluorootic acid (FOA) and resistance to 200 μg/ml G418, respectively^51^. The cassette was subsequently replaced with a gene expression module consisting of a natural *S. cerevisiae* promoter (either P_SIR2_, P_RAP1_, or P_SUP35_), a fusion protein-encoding gene, and a terminator (either t_TEF_, t_ADH1_, or t_CYC1_). Conventional integration of pRS304-encoding Sir3-SET was also employed^52^.

Gene deletions were performed by transforming cells with DNA fragments – containing sequence homologous to target loci at their ends – encoding *KanMX* (conferring resistance to 200 μg/ml G418), *NatMX* (conferring resistance to 100 μg/ml nourseothricin sulfate), *HphMX* (conferring resistance to 300 μg/ml hygromycin B), *HIS3^Sk^MX* (conferring histidine prototrophy), or *LYS2* (conferring lysine prototrophy). All chromosomal genetic modifications were confirmed by PCR analysis of both sites of homologous recombination. Non-chromosomally encoded proteins were produced from the *ARS CEN* plasmid pRS315^52^, derivatives of which were maintained in cells grown in medium lacking leucine.

Plate-based phenotypic analyses were performed by normalizing overnight cell cultures to an OD600 of 5.0, serially diluting cells in phosphate-buffered saline (PBS), and spotting dilutions onto medium containing limiting adenine (10 μg/ml; Ade^10^) and/or 1 mg/ml FOA in the presence or absence of aTc. Plates were incubated for three days at 30°C and stored at 4°C for two days before being photographed.

*S. cerevisiae* strains and plasmids used in this study are listed in Table S1.

### Calculating the probability of silent information loss

Round 1 colonies grown under adenine-replete conditions without selection were thoroughly resuspended in PBS and diluted. ∼250 colony forming units (CFUs) were plated on either selective medium containing FOA and Ade^10^ or non-selective, adenine-replete medium lacking FOA. The fraction of silent information-retaining cells was determined by dividing the number of pigmented, FOA-resistant colonies by the number of CFUs plated.

We calculated the probability of silent information loss per generation (*P*) in a population of cells undergoing exponential growth using the following equation: *P*^N^ = *x*, where N equals the number of generations that occurred during the formation of each Round 1 colony tested, and where *x* equals the fraction of FOA-resistant cells per colony. This calculation assumes that differences in the growth rate and replicative lifespan of cells carrying either silent or active reporter genes are negligible during a round of colony formation.

### Chromatin immunoprecipitation and quantitative PCR

50-100 ml cultures of *S. cerevisiae* cells were grown in YPAD medium to an OD_600_ of 1 to 2. Cells were fixed in the presence of 1% formaldehyde for 15 minutes at room temperature with gentle mixing on a moving platform. For Sir3-FLAG_3X_-SET immunoprecipitations, cells were fixed in the presence of 1.5 mM ethylene glycol bis(succinimidyl succinate) (EGS) for 30 minutes at room temperature and then fixed in the presence of 1% formaldehyde for 30 minutes at room temperature. Crosslinking reactions were quenched in the presence of 125 mM glycine. Crosslinked cells were pelleted, washed twice in cold tris-buffered saline (TBS), once in cold water, and stored at -80°C.

Frozen samples were thawed and resuspended in cold ChIP lysis buffer (0.1% sodium dodecyl sulfate, 1% Triton X-100, 0.1% sodium deoxycholate, 140 mM NaCl, 1 mM EDTA, 50 mM HEPES-KOH pH 7.5) supplemented with commercial protease inhibitor tablets. Cells were disrupted in a cold room by bead beating for a total of six minutes, with one-minute incubations in an ice-cold water bath following every minute of disruption. The insoluble, chromatin-containing fractions of broken cell lysates were collected by centrifugation in a cooling centrifuge set at maximum velocity (4°C at 21,130 g) for five minutes. The insoluble fractions were washed once with 1 ml of ChIP lysis buffer, pelleted again by centrifugation, and thoroughly resuspended in 600 μl of ChIP lysis buffer. Resuspended samples were sonicated in a cold room using a tip sonicator (Sonics VCX 500; set at 30% amplitude) for a total of 80 seconds, with five-minute incubations on ice after every 20 seconds of sonication. The soluble, chromatin-containing fractions of sonicated samples were collected by centrifugation in a cooling centrifuge set at maximum velocity for 15 minutes.

The protein concentration of chromatin samples was measured by the Bradford method against a bovine serum albumin standard curve. 600 μg of chromatin was precleared with 30 μl of Protein A- or Protein G-coupled magnetic beads (Invitrogen Dynabeads) for one hour at 4°C on a rotating platform. For each immunoprecipitation, 100-200 μg of precleared chromatin was incubated at 4°C for three hours with magnetic beads that had been prebound to one of the following antibodies: 3 μg of α-V5 antibody (Abcam ab9116), 1.5 μg of α-H3K9me2 antibody (Abcam ab1220), 3 μg of α-H3K9me3 antibody (Abcam ab8898), 4 μg of α-FLAG antibody (Sigma M2), or 4 μg of α-RPB1 (Biolegend 8WG16).

Immunoprecipitations and quantitative PCR (qPCR) analysis of immunoprecipitates were performed essentially as described^53^, with the following modifications: samples were washed three times in ChIP lysis buffer, DNA was purified using an E.Z.N.A. Cycle Pure Kit (Omega Bio-tek), and real-time PCR was performed using the QuantStudio 6 Pro system (Applied Biosystems). Primers used for qPCR analysis were designed in Primers3 4.1.0 and are listed in Table S2.

### Chromatin immunoprecipitate sequencing

Chromatin immunoprecipitates and input DNA samples were further sheared by sonication using a Q800R3 sonicator (Qsonica; set at 20% amplitude). Samples were sonicated for a total of 15 minutes, with 15-second incubations without sonication following every 15 seconds of sonication. DNA was purified using a NucleoSpin Gel and PCR Clean-up kit (Machery-Nagel). Samples were barcoded and pooled as previously described^54^. Pooled samples were sequenced on an Illumina NextSeq 500 system at the Bauer Core Facility (Harvard University). In-line barcode demultiplexing was performed courtesy of the Bauer Core Facility, and reads were mapped to the S288C reference genome R64.3.1, the sequence of which was manually edited to lack the native S288C *ADE2* gene and to contain *tetO*_10X_-*ADE2* or *URA3^Kl^*- *tetO*_10X_-*ADE2* reporters in place of either *HMR* or *HFL1*. Read alignments were performed using Bowtie with default settings. Data was normalized to counts per million using the bamCoverage module of deepTools 3.5.2 and visualized in IGV 2.16.0.

### RNA extraction and cDNA synthesis

Total RNA was collected from *S. cerevisiae* cells by hot acid phenol-chloroform extraction. RNA samples were treated with TURBO DNase (Invitrogen) and purified using a Quick-RNA Miniprep Kit (Zymo Research). cDNA was synthesized using the SuperScript IV First-Strand Synthesis System (Invitrogen) with Oligo(dT) primers. cDNA was diluted 1:10 or 1:50, and cDNA levels were quantified by real-time PCR as previously described^53^. Primers used for qPCR analysis were designed in Primers3 4.1.0 and are listed in Table S2.

### Western blot analysis

Yeast whole cell extracts were prepared by bead beating and trichloroacetic acid (TCA)- mediated protein precipitation. Samples were solubilized in sodium dodecyl sulfate (SDS) loading buffer containing 50 mM Cleland’s reagent (DTT), boiled for five minutes, and clarified by centrifugation. Gel electrophoresis was performed using NuPAGE 4-12% Bis-Tris precast gels (Invitrogen) and MOPS-SDS running buffer (Boston Bioproducts). Samples were transferred to nitrocellulose membranes, which were stained with Ponceau S Staining Solution (Thermo Scientific), photographed, and blocked in tris-buffered saline containing 0.1% Tween 20 (TBST) and 5% milk. The same antibodies used for chromatin immunoprecipitations were also used to probe membranes for epitope-tagged proteins of interest. Primary antibodies were diluted 1:2,500 and secondary antibodies conjugated to HRP were diluted 1:5,000 in TBST milk. Chemilluminescence was detected by a ChemiDoc XRS+ System (Bio-rad).

### Transposon mutagenesis and mapping of transposition sites

*hmr*Δ::*tetO*_10X_-*ADE2* reporter-containing *sir*^−^ cells producing chromosomally encoded TetR-SET, Sir2*^Sp^*-HP1, and Sir3-SET were transformed with *ARS CEN* plasmid pSG36, which encodes *URA3*, a *Hermes-NatMX* transposon, and a hyperactive Hermes transposase variant produced under the control of a galactose-inducible promoter^55^. Six transformants were individually cultured in glucose-containing medium lacking uracil. Cell cultures were diluted to an OD_600_ of 0.05 in 25 ml of galactose-containing medium lacking uracil − to induce transposition − and grown to saturation overnight. The next day, cell cultures were diluted to an OD_600_ of 0.25 in 100 ml of glucose-, uracil-, and FOA-containing medium – to cure cells of pSG36 – and grown to saturation overnight. The following day, cell cultures were diluted to an OD600 of 0.5 in 100 ml of glucose-, uracil-, FOA-, and nourseothricin sulfate-containing medium – to select for transposon-containing cells lacking pSG36 – and grown to saturation overnight. Finally, cells were plated on glucose-, uracil-, FOA-, nourseothricin sulfate-, and 10 μg/ml adenine (Ade^10^)-containing medium in the presence or absence of aTc. Plates were incubated for three days at 30°C and stored at 4°C for two days before being visually inspected.

Genomic DNA of mutants exhibiting altered colony pigmentation was purified, and *Hermes-NatMX* transposon insertion sites were mapped by arbitrary PCR^56^ and Sanger sequencing. Primers used for arbitrary PCR are listed in Table S2.

### Data availability

Raw and processed ChIP-seq data is available at the NCBI Gene Expression Omnibus (accession number: GSE236173).

---

**Figure S1.**
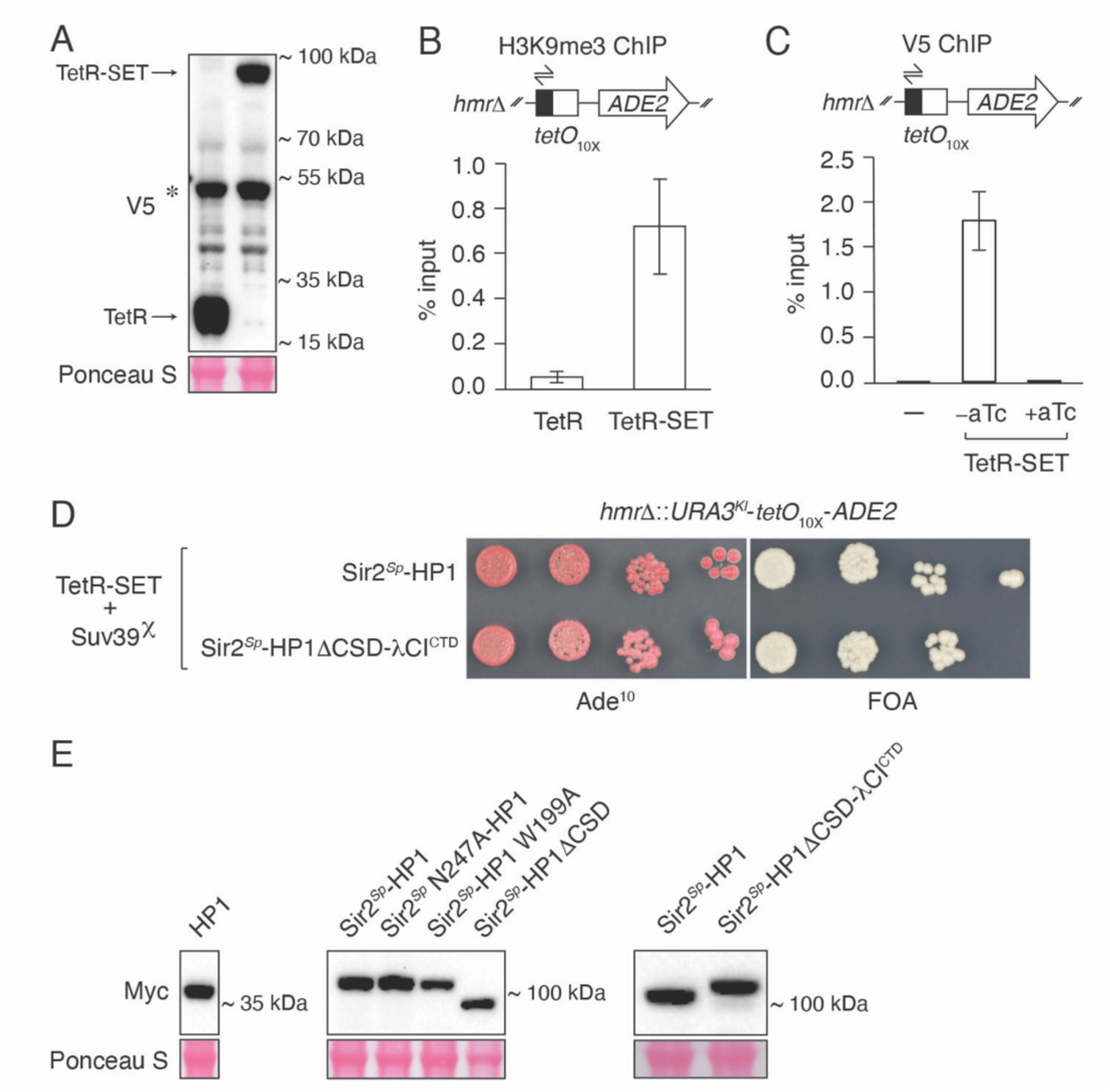
TetR-SET and Sir2*^Sp^*-HP1 repress *tetO*_10X_-*ADE2* reporter gene expression. **(A)** Western blot analysis of whole cell extracts prepared from cells producing V5 epitope-tagged TetR or TetR-SET. Arrows point to bands corresponding to these proteins. The asterisk indicates a non-specific band. Ponceau S membrane staining serves as a loading control. **(B)** Trimethylated H3K9 (H3K9me3) levels detected by chromatin immunoprecipitation and quantitative PCR (ChIP-qPCR) in *tetO*_10X_-*ADE2* reporter-containing cells producing TetR or TetR-SET using primers (half-headed arrows) targeting unique DNA sequence (black rectangle) present adjacent to *tetO*_10X_ (white square). Values represent the averages and standard deviations of three biological replicates. **(C)** ChIP-qPCR analysis of TetR-SET levels at *tetO*_10X_. Cells either lacked or produced TetR- SET, which contains a V5 epitope tag. TetR-SET-containing cells were either cultured in the absence of aTc or in the presence of aTc for 30 minutes. Values represent averages and standard deviations of three biological replicates. The level of TetR-SET at *tetO*_10X_ in aTc-treated cells approached the limit of detection. **(D)** Phenotypes of cells producing either Sir2*^Sp^*-HP1 or a Sir2*^Sp^*-HP1 variant containing the C-terminal dimerization domain of bacteriophage λ CI in place of the Chp2 chromo shadow domain (Sir2*^Sp^*-HP1ΔCSD-λCI^CTD^). In addition to TetR-SET, these cells produce an H3K9me “read-write” protein, which we refer to as Suv39^χ^, and contain a second reporter gene, *URA3^Kl^*, positioned upstream of *tetO*_10X_-*ADE2*. Sir2*^Sp^*-HP1 and Sir2*^Sp^*-HP1ΔCSD-λCI^CTD^ were encoded on pRS315. Cells were spotted on leucine-lacking medium containing limiting adenine (10 μg/ml; Ade^10^) or 5-fluoroorotic acid (FOA). **(E)** Western blot analysis of whole cell extracts prepared from cells producing Myc_3X_ epitope-tagged HP1, Sir2*^Sp^*-HP1, or the indicated Sir2*^Sp^*-HP1 variant. Ponceau S membrane staining serves as a loading control.

**Figure S2.**
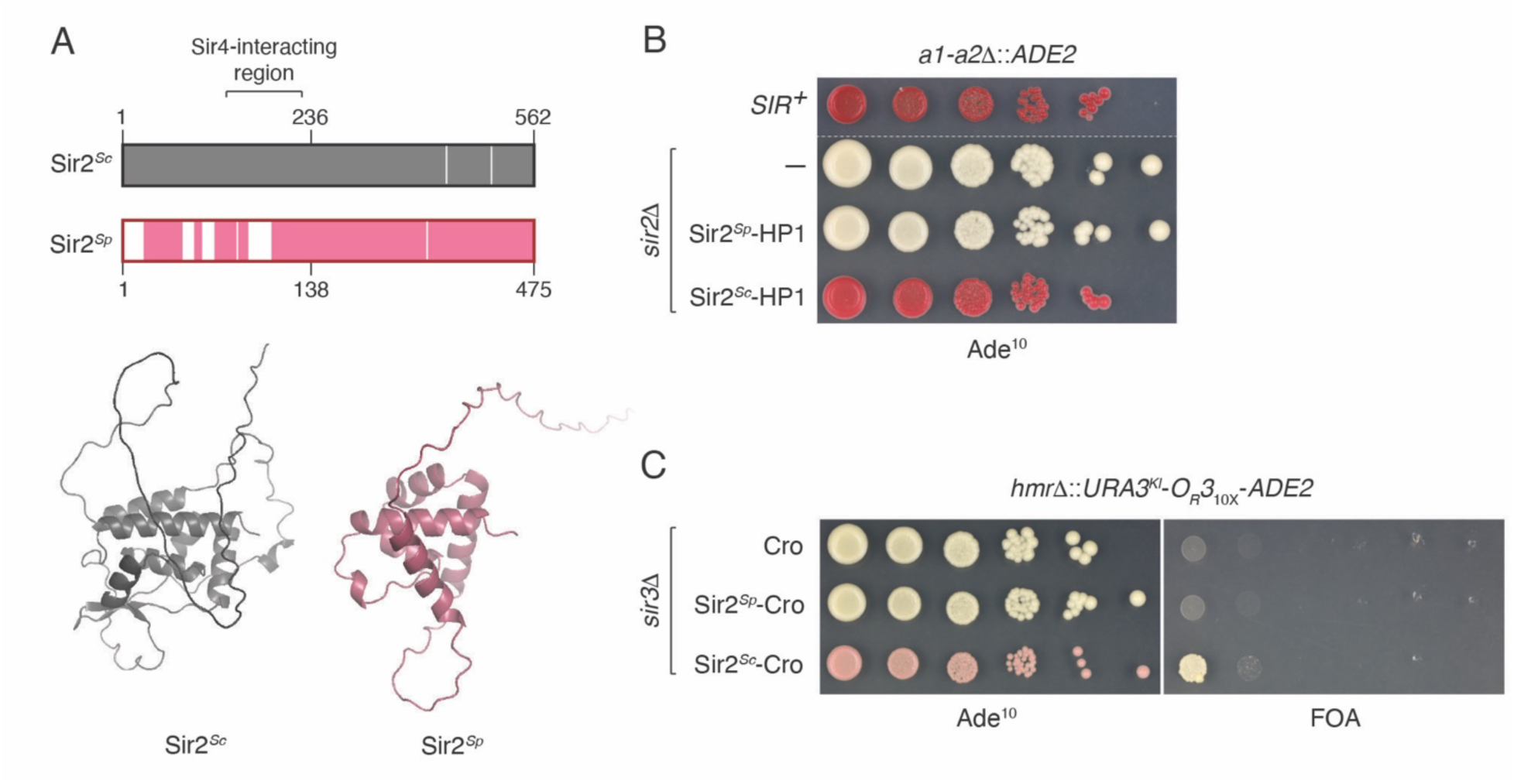
Sir2*^Sp^*-HP1 functions independently of native *S. cerevisiae* silencing factors. **(A)** Clustal W alignment of Sir2*^Sc^* (gray) with Sir2*^Sp^*(pink). Regions lacking sequence conservation are indicated by colorless gaps. AlphaFold predicted structures of the Sir2*^Sc^* N-terminal domain (residues 1-236 of Sir2*^Sc^*; gray) and the Sir2*^Sp^*N-terminal domain (residues 1- 138 of Sir2*^Sp^*; raspberry) are shown below for comparison. **(B)** Phenotypes of *SIR^+^* cells (producing the native SIR complex) and *sir2*Δ cells containing an *ADE2* reporter gene situated between the natural *E* and *I* silencers of *HMR* in place of *a2* and *a1*. Sir2*^Sp^*-HP1 and Sir2*^Sc^*-HP1 were encoded on pRS315. Cells were spotted on Ade^10^ medium lacking leucine. The white dashed line indicates a site of cropping between spots of *SIR^+^* cells grown and photographed on a separate plate. **(C)** Phenotypes of cells producing bacteriophage λ Cro, a Sir2*^Sp^*-Cro fusion protein, or a Sir2*^Sc^*- Cro fusion protein. Cells contain a modified genetic reporter featuring ten tandem λ operators (*O_R_3*_10X_) situated between *URA3^Kl^*and *ADE2*. Cro and Sir2-Cro fusion proteins were encoded on pRS315. Cells were spotted on leucine-lacking medium containing Ade^10^ or FOA.

**Figure S3.**
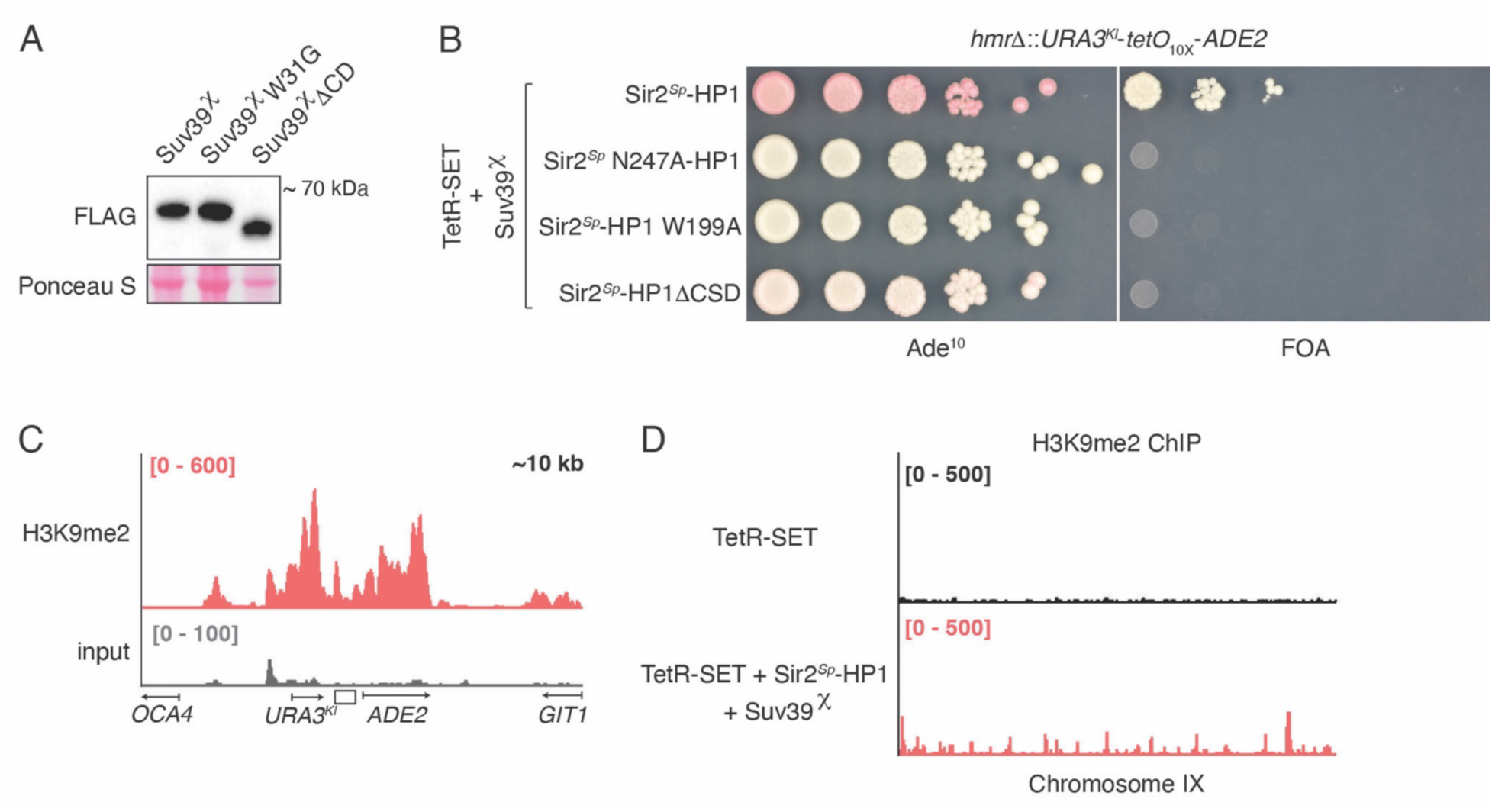
Suv39^χ^-mediated read-write promotes spreading of H3K9me but also leads to off-target H3K9me. **(A)** Western blot analysis of whole cell extracts prepared from cells producing FLAG_3X_ epitope-tagged Suv39^χ^, a Suv39^χ^ variant containing a mutant chromodomain incapable of methyllysine recognition (Suv39^χ^ W31G), or a Suv39^χ^ variant lacking its chromodomain (Suv39^χ^ΔCD). Ponceau S membrane staining serves as a loading control. **(B)** Phenotypes of *URA^Kl^*-*tetO*_10X_-*ADE2* reporter-containing cells producing TetR-SET, Suv39^χ^, and either Sir2*^Sp^*-HP1, a Sir2*^Sp^*-HP1 variant lacking histone deacetylase activity (Sir2*^Sp^* N247A- HP1), a Sir2*^Sp^*-HP1 variant containing a mutant chromodomain incapable of methyllysine recognition (Sir2*^Sp^*-HP1 W199A), or a monomeric Sir2*^Sp^*-HP1 variant lacking the Chp2 chromo shadow domain (Sir2*^Sp^*-HP1ΔCSD). Sir2*^Sp^*-HP1 and Sir2*^Sp^*-HP1 variants were encoded on pRS315. Cells were spotted on leucine-lacking medium containing Ade^10^ or FOA. **(C)** Chromatin immunoprecipitate sequencing (ChIP-seq) profile of H3K9me2 at the *URA^Kl^*- *tetO*_10X_-*ADE2* reporter in *sir3*Δ cells producing chromosomally encoded TetR-SET, Sir2*^Sp^*-HP1, and Suv39^χ^. Data ranges (in counts per million) are indicated in brackets. The region visualized encompasses ∼10 kb of chromosome III centered about *tetO*_10X_. The H3K9me2 profile is shown in red, and the input DNA profile is shown in gray. The position of *tetO*_10X_ is indicated by a white rectangle, and select genes are indicated by arrows. **(D)** ChIP-seq profiles of H3K9me2 in reporter-containing *sir3*Δ cells either producing TetR-SET alone (black track) or producing TetR-SET, Sir2*^Sp^*-HP1, and Suv39^χ^ (red track). Data ranges (in counts per million) are indicated in brackets. The region visualized is chromosome IX.

**Figure S4.**
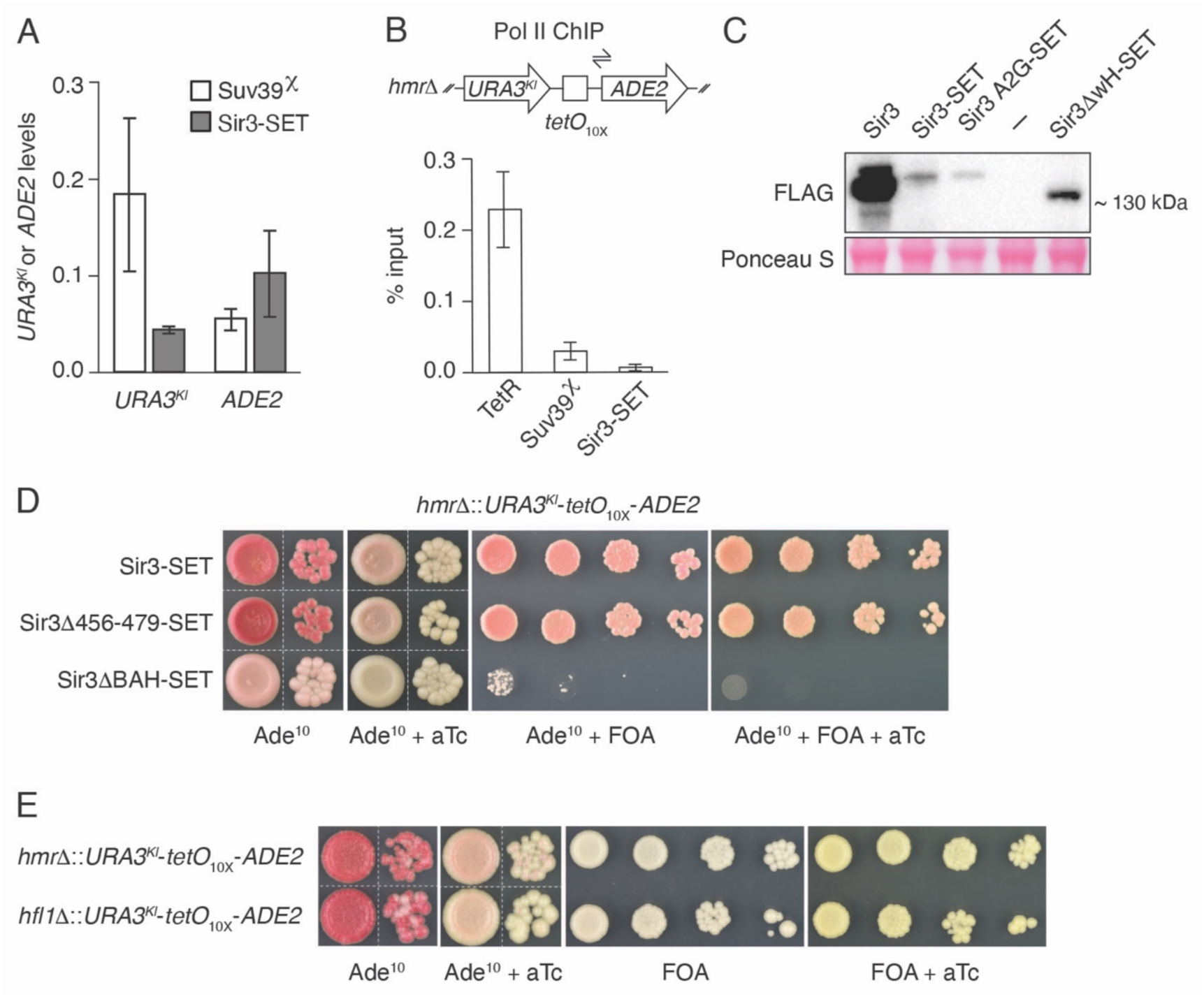
Heritable reporter gene silencing occurs independently of specific DNA sequence. **(A)** Reverse transcriptase quantitative PCR (RT-qPCR) analysis of *URA3^Kl^*and *ADE2* mRNA levels in two strains: a *sir3*Δ strain producing TetR-SET, Sir2*^Sp^*-HP1, and Suv39^χ^ (white), and a *sir*^−^ strain producing TetR-SET, Sir2*^Sp^*-HP1, and Sir3-SET (gray). Values represent *URA3^Kl^* and *ADE2* levels (normalized to *ACT1*) in cells of the indicated strain relative to *URA3^Kl^* and *ADE2* levels in unsilenced reporter gene-containing *sir3*Δ cells producing TetR alone. Averages and standard deviations of three biological replicates are shown. **(B)** ChIP-qPCR analysis of Rpb1 – the large subunit of RNA polymerase II (Pol II) – levels at the singular *ADE2* promoter in *sir3*Δ cells producing TetR alone (left bar), *sir3*Δ cells producing TetR-SET, Sir2*^Sp^*-HP1, and Sir3-SET (middle bar), or *sir*^−^ cells producing TetR-SET, Sir2*^Sp^*- HP1, and Sir3-SET (right bar). Primers targeting the promoter region of *ADE2* are shown (half- headed arrows). Values represent the averages and standard deviations of three biological replicates. **(C)** Western blot analysis of whole cell extracts prepared from cells containing FLAG_3X_ epitope-tagged Sir3, Sir3-SET, a Sir3-SET variant harboring a mutation that prevents Sir3 acetylation and disrupts bromo adjacent homology domain structure (Sir3 A2G-SET), or a monomeric Sir3- SET variant lacking its winged helix (wH) domain (Sir3ΔwH-SET). All proteins were encoded on pRS315 and produced under the control of the *SUP35* promoter. Ponceau S membrane staining serves as a loading control. **(D)** Colony color and FOA sensitivity phenotypes of *sir*^−^ cells containing chromosomally encoded TetR-SET and Sir2*^Sp^*-HP1 as well as pRS315-encoded Sir3-SET, a Sir3-SET variant lacking Rap1 interaction determinants (Sir3Δ456-479-SET), or a Sir3-SET variant lacking its bromo adjacent homology (BAH) domain (Sir3ΔBAH-SET). Cells were spotted on leucine-lacking medium containing the indicated compounds. White dashed lines indicate sites of cropping between spots of cells grown and photographed on the same plate. **(E)** Phenotypes of *sir*^−^ cells containing a *URA3^Kl^*-*tetO*_10X_-*ADE2* reporter in place of either *HMR* or *HFL1*. Cells also contain chromosomally encoded TetR-SET, Sir2*^Sp^*-HP1, and Sir3-SET. Cells were spotted on medium containing the indicated compounds. White dashed lines indicate sites of cropping between spots of cells grown and photographed on the same plate.

**Figure S5.**
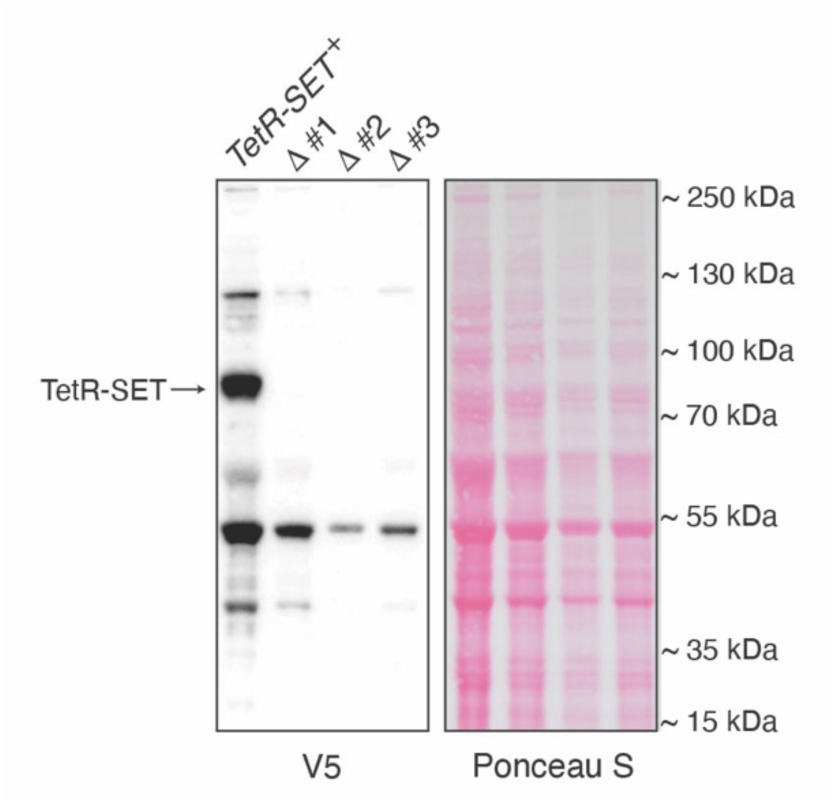
Reporter gene silencing can be maintained in cells devoid of TetR-SET. Western blot analysis of whole cell extracts prepared from *sir*^−^ cells that produce Sir2*^Sp^*-HP1 and Sir3-SET but either contain or lack the gene encoding V5 epitope-tagged TetR-SET. The arrow points to the band corresponding to TetR-SET. Ponceau S membrane staining serves as a loading control. Three deletion-containing strains (Δ #1-3) carry the *Saccharomyces kluyveri HIS3* gene (*HIS3^Sk^*) in place of the TetR-SET-encoding gene and were isolated on FOA-containing medium lacking histidine.

**Figure S6.**
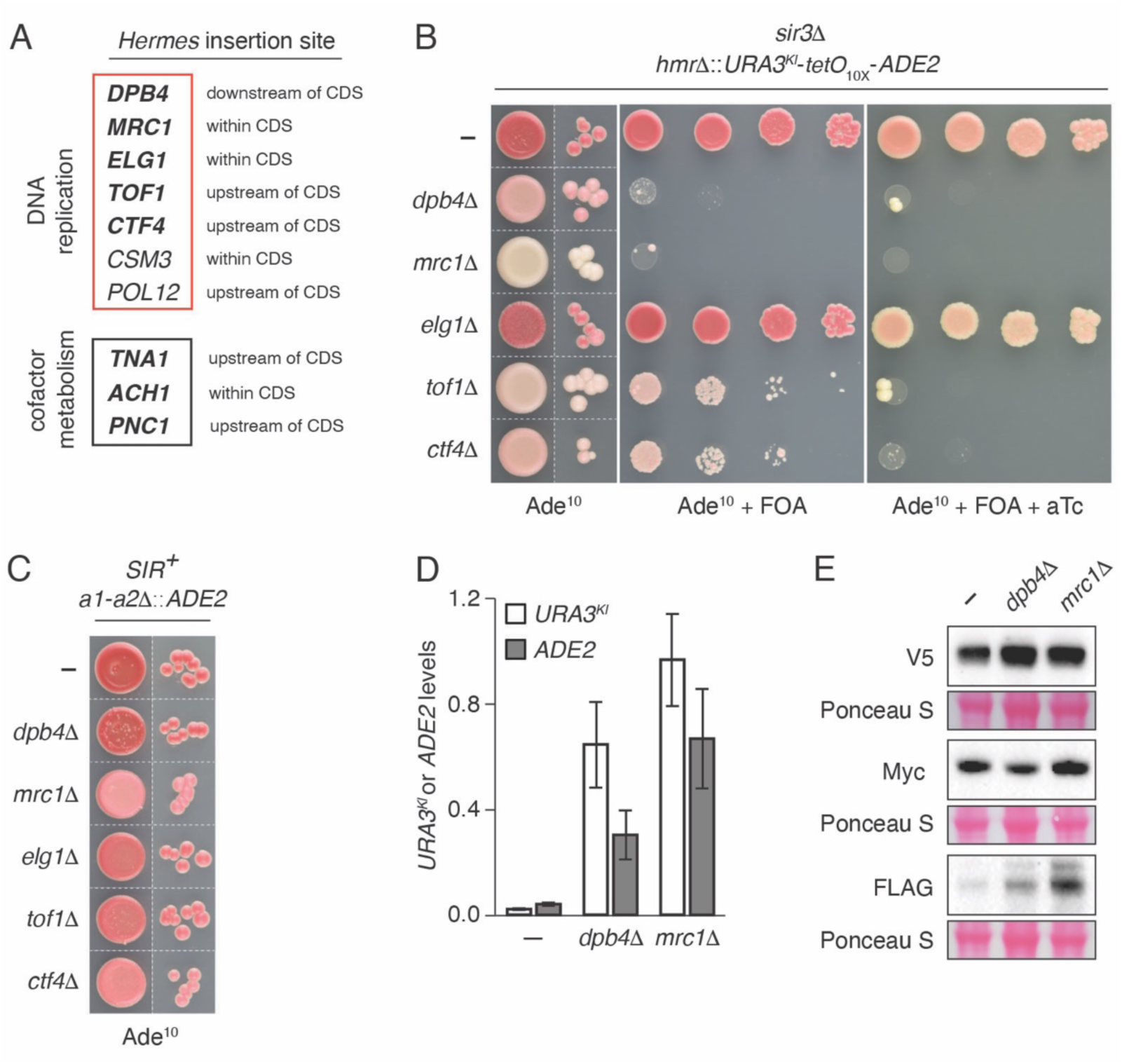
Reporter gene silencing is sensitive to loss of chromatin replication factors. **(A)** List of *Hermes* transposon insertions that impair *URA^Kl^*-*tetO*_10X_-*ADE2* reporter gene silencing in *sir*^−^ cells producing TetR-SET, Sir2*^Sp^*-HP1, and Sir3-SET. Genes predicted to be affected by transposition are grouped by their involvement in either DNA replication (red box) or cofactor metabolism (black box). Sites of transposition – relative to the coding sequence (CDS) of each gene – are described. Genes highlighted in bold were identified at least twice and up to five times, whereas *CSM3* and *POL12* were identified once. **(B)** Phenotypes of *URA3^Kl^*-*tetO*_10X_-*ADE2* reporter gene-containing cells lacking one of five replisome-associated genes. These cells are *sir3*Δ and contain chromosomally encoded TetR-SET, Sir2*^Sp^*-HP1, and Sir3-SET. Cells were spotted on medium containing the indicated compounds. **(C)** Phenotypes of *SIR^+^* cells lacking one of five replisome-associated genes. Cells produce the native SIR complex and contain an *ADE2* reporter gene situated between the natural *E* and *I* silencers of *HMR*. **(D)** RT-qPCR analysis of *URA3^Kl^*(white) and *ADE2* (gray) mRNA levels in *URA^Kl^*-*tetO*_10X_- *ADE2* reporter gene-containing *sir3*Δ cells producing TetR-SET, Sir2*^Sp^*-HP1, and Sir3-SET. Cells either lacked or contained one of two mutations disrupting replisome function (*dpb4*Δ or *mrc1*Δ). Values represent *URA3^Kl^* and *ADE2* levels (normalized to *ACT1*) in these cells relative to *URA3^Kl^* and *ADE2* levels in reporter gene-containing *sir3*Δ cells producing TetR alone. Averages and standard deviations of three biological replicates are shown. **(E)** Western blot analysis of whole cell extracts prepared from *dpb4*Δ and *mrc1*Δ cells (or cells containing a fully functional replisome) producing TetR-SET, Sir2*^Sp^*-HP1, and Sir3-SET. TetR- SET contains a V5 epitope tag, Sir2*^Sp^*-HP1 contains a Myc_3X_ epitope tag, and Sir3-SET contains a FLAG_3X_ epitope tag. Ponceau S membrane staining serves as a loading control.

**Table S1.**
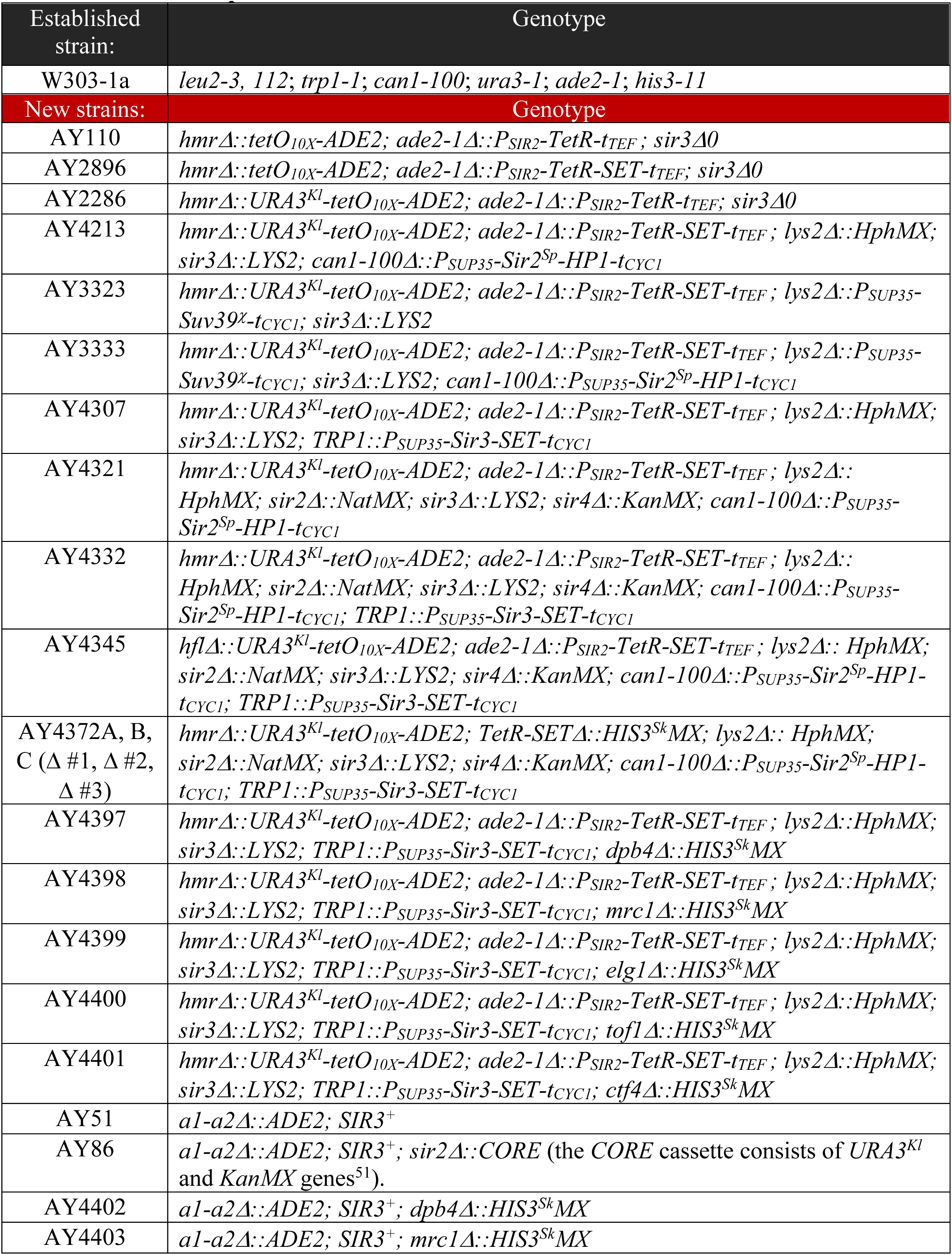
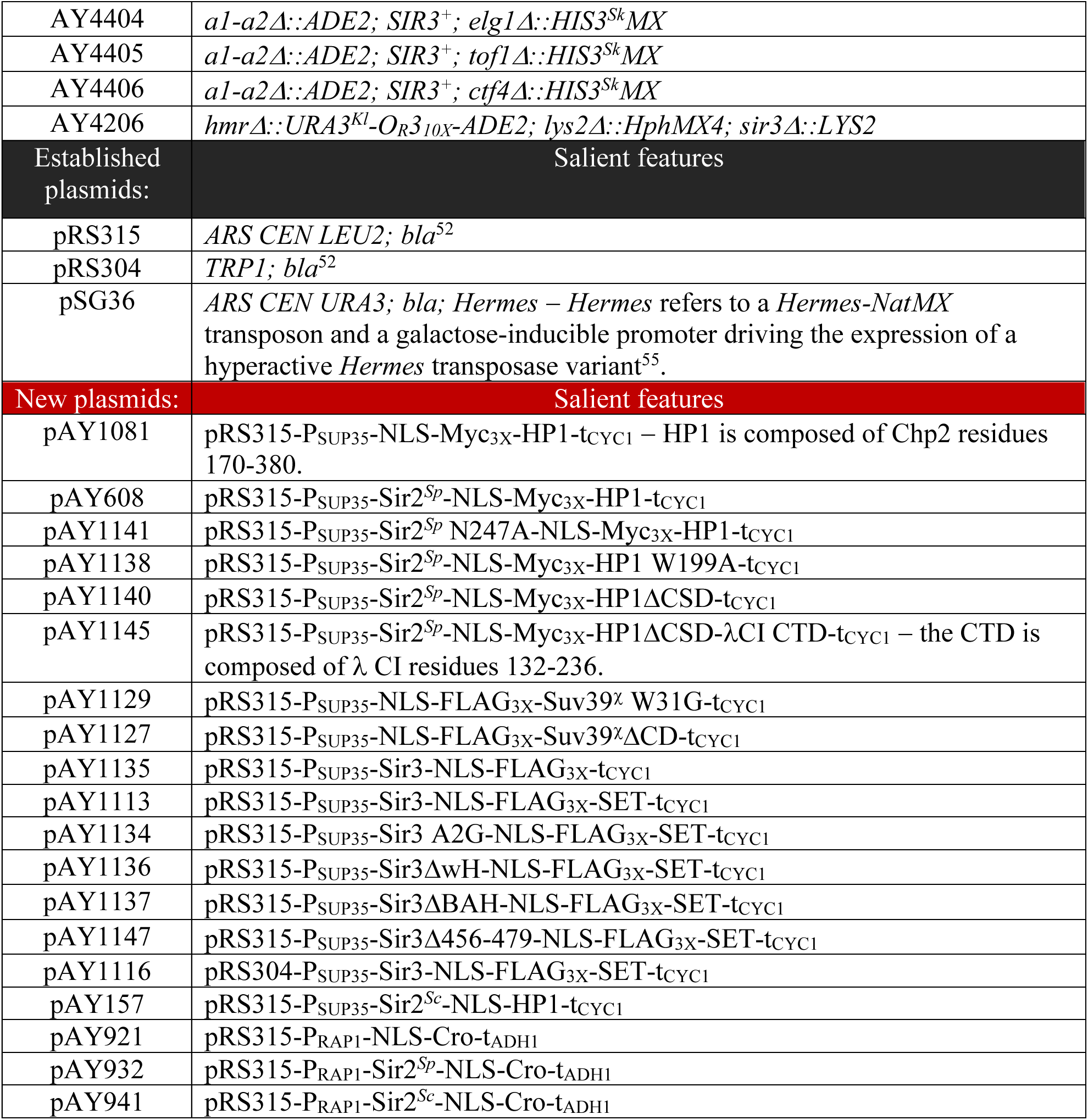
Strains and plasmids.

**Table S2.**
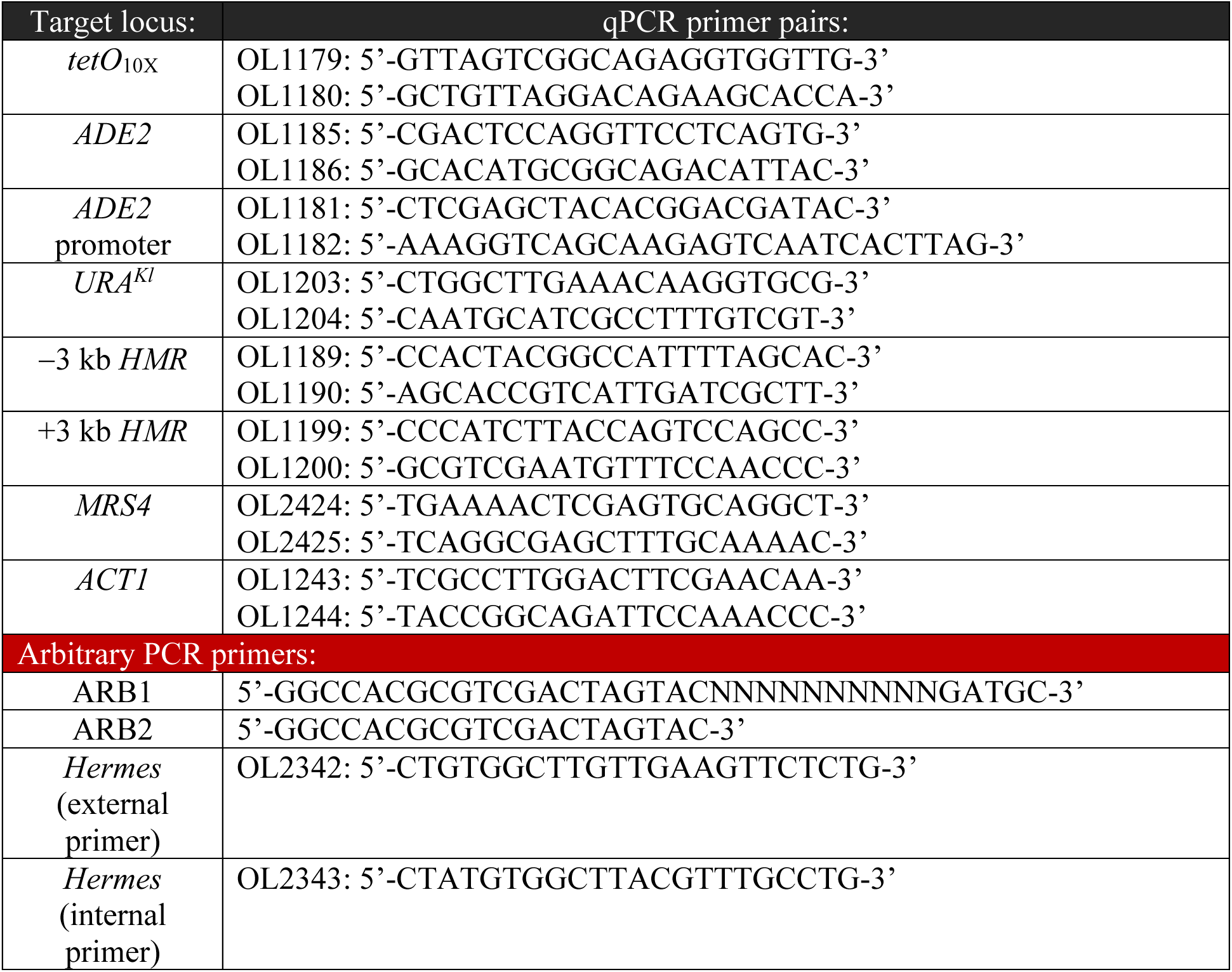
Primers.

